# Distinct Binding Modes of Inter-Spike Cross-Linking Suggest a Supplementary Mechanism for SARS-CoV-2 Antibody Neutralization

**DOI:** 10.1101/2024.11.07.622402

**Authors:** Xuanyu Nan, Yujie Li, Rui Zhang, Ruoke Wang, Niannian Lv, Jiayi Li, Yuanfang Chen, Bini Zhou, Yangjunqi Wang, Ziyi Wang, Jiayi Zhu, Jing Chen, Jinqian Li, Wenlong Chen, Qi Zhang, Xuanling Shi, Changwen Zhao, Chunying Chen, Zhihua Liu, Yuliang Zhao, Dongsheng Liu, Xinquan Wang, Li-Tang Yan, Taisheng Li, LinQi Zhang, Yuhe R. Yang

## Abstract

The emergence of severe acute respiratory syndrome coronavirus 2 (SARS-CoV-2) and its Omicron subvariants drastically amplifies transmissibility, infectivity, and immune escape, mainly due to their resistance to most neutralizing antibodies. Thus, exploring the mechanisms underlying antibody evasion is crucial. Although the full-length native form of antibody, immunoglobulin G (IgG), offers valuable insights into the neutralization, structural investigations primarily focus on the fragment of antigen-binding (Fab). Here, we employ single-particle cryo-electron microscopy (cryo-EM) to characterize a W328-6H2 antibody, in its native IgG form complexed with severe acute respiratory syndrome (SARS), severe acute respiratory syndrome coronavirus 2 wild-type (WT) and Omicron variant BA.1 spike protein (S). Three high-resolution structures reveal that the full-length IgG forms a centered head-to-head dimer of trimer when bound fully stoichiometrically with both SARS and WT S, while adopting a distinct offset configuration with Omicron BA.1 S. Combined with functional assays, our results suggest that, beyond the binding affinity between the RBD epitope and Fab, the higher-order architectures of S trimer and full-length IgG play an additional role in neutralization, elucidate a potential molecular basis of Omicron immune escape, and expand our understanding on antibody evasion mechanisms.

## Main Text

### INTRODUCTION

Severe acute respiratory syndrome coronavirus 2 (SARS-CoV-2) has been continuously evolving through mutations in its viral genome with increased transmissibility, infectivity, and immune evasion^1, 2^. In particular, the latest variant of concern (VOC), Omicron, is distinguished by having the largest genetic divergence from the wild-type (WT) virus, and its sublineages including BA.1, BA.2, BA.4, BA.5, BF.7 and XBB have become the most prevalent SARS-CoV-2 variants globally. With a total of over 30 amino acid substitutions in spike (S) including 15 in the receptor binding domain (RBD) (G339D, S371L, S373P, S375F, K417N, N440K, G446S, S477N, T478K, E484A, Q493R, G496S, Q498R, N501Y, and Y505H), the magnitude of polyclonal antibody evasion by BA.1 is more pronounced than WT after receiving primary vaccination or infection^3^. Additionally, most SARS-CoV-2 specific monoclonal antibodies (mAbs), including those approved by the FDA, exhibit partial or near-complete ineffectiveness against the Omicron variants^4, 5, 6, 7^.

With the capability of high-throughput screening, mAbs have been categorized into different epitope groups, mainly RBD and N-terminal domain (NTD). mAbs targeting the most immunogenic RBD sites have been clustered by a combination of competitive binding assays and electron microscopy (EM)-based epitope mapping^4, 5, 8, 9, 10^. Prior to the few recent studies^11, 12, 13^, structural characterization primarily focused on assessing single-Fab footprints when evaluating potency, breadth, or determining mechanisms of neutralization. However, the extent to which Fab can fully account for all antibody functions has been a topic of ongoing debate due to the fact that IgGs often display stronger binding avidity and higher neutralizing potency compared to Fabs^14, 15, 16, 17^. Recently, it was reported that certain full-length IgGs targeting the ACE2 site could induce a higher number of “up” RBD through their bivalent binding, leading to enhanced S1 subunit shedding and improved antiviral potency^11^. Furthermore, researchers also demonstrated that both the WT^13^ and BA.1^12^ can form trimer-dimer structures upon antibody binding, and hypothesized that the neutralization mechanism involved virion aggregation due to IgG cross-linking. More importantly, the Y-shaped structure is considered to be more biologically relevant, as it is preserved in both the membrane-bound Immunoglobulin (mIg) form (B cell receptor, BCR)^18, 19, 20^ during antibody affinity maturation and in the secreted Immunoglobulin (sIg) form (mAb). IgG not only provides information about the epitope of a single-Fab arm, but also exhibits a bivalent binding mode involving both Fab arms. Therefore, it is crucial to explore the underlying mechanisms of antibodies from IgG-related structures that are closely correlated to functional capabilities.

With a substantial portion of structural analysis focused on epitope interactions of Fab-RBD/ S trimer complexes, it is generally recognized that neutralization evasion of the Omicron variant is highly relevant with binding affinity, resulting from epitope mutations^1^. At the same time, studies on antibody/antigen binding employing multimeric constructs suggest that avidity^11, 21^ and occupancy^22^ also impact the neutralization of monoclonal antibodies. Nevertheless, the neutralization mechanisms of antibodies have long been intricate and sophisticated^23^, with a full understanding of the precise mechanisms of action for many neutralizing antibodies still unfolding. A wide array of factors influencing these mechanisms awaits thorough investigation.

In this work, we integrated negative-stain electron microscopy (ns-EM) and cryo-EM structural analysis with functional assays including flow cytometry (FC), surface plasmon resonance (SPR), pseudovirus (PsV) neutralization assays, and bead-based aggregation assays, to characterize an antibody named W328-6H2 (6H2). This antibody, isolated from a severe acute respiratory syndrome (SARS) patient^24^, was analyzed in its native full-length IgG form bound to trimeric sarbecovirus S proteins, encompassing SARS, WT, and Omicron variant BA.1. By correlating binding avidity and neutralization profiles with cross-linking patterns, we presented a potential mechanism for antibody neutralization differences through inter-spike cross-linking. Our findings highlight the additional role of multivalent binding in enhancing avidity and neutralization potency of 6H2 IgG against SARS and WT and provide new insights into how different higher-order structures, beyond epitope mutations, contribute to the loss of neutralization against Omicron variants.

## RESULTS

### Distinct dimer-trimer architectures of IgG -WT/ Omicron S complexes

As detailed in our recent published work^24^, a total of 60 RBD-targeting mAbs were isolated from SARS patients. Seven major groups (RBD-1 to RBD-7) were identified through a combination of SPR competition assay and ns-EM analysis. Among them, RBD-5 antibodies, targeting the outer face of RBD, displayed the broadest cross-neutralization towards SARS and WT. In RBD-5 class, 6H2 IgG exhibited the most potent neutralization against WT, however, its neutralization was entirely diminished against the BA.1. To further investigate the cross-neutralization of 6H2 among SARS-CoV-2 variants, we applied PsV neutralization and cell surface stain assays against a panel of major SARS-CoV-2 variants from BA.1 to CH.1.1 (Fig. 1a, Supplementary Fig.1 and 2). Our initial observations revealed that binding ability of 6H2 IgG with cell surface expressed SARS-CoV-2 variant S did not align with trends in neutralization potencies. Specifically, the binding ability of 6H2 IgG to BA.1 showed only a slight reduction compared to other variants: 2.4-fold for WT, 2.3-fold for Alpha, 2.6-fold for Beta, 1.3-fold for Gamma, and 3-fold for Delta (Fig. 1a). Despite this, neutralization against BA.1 was completely abolished (Fig. 1a). In contrast, analyses focusing solely on the Fab component showed that binding ability of 6H2 Fab with cell surface expressed SARS-CoV-2 variant S (TFI) aligned perfectly with neutralization potencies (IC50). The data displayed a decrease from 29.73 to 3.07 when comparing SARS to WT, remaining relatively stable up to the Gamma variant before plummeting dramatically to 0.09 for BA.1 (Supplementary Fig.1). Similarly, neutralization potency, IC50 values, increased from 0.15 to 0.88 when comparing SARS to WT, stabilized around 1 up to Gamma, and then dramatically increased to >100 for BA.1(Supplementary Fig.1). These discrepancies between 6H2 IgG binding and neutralization prompted us to explore other factors that could contribute to the complete loss of neutralization of 6H2 IgG against the Omicron variants.

**Fig. 1.**
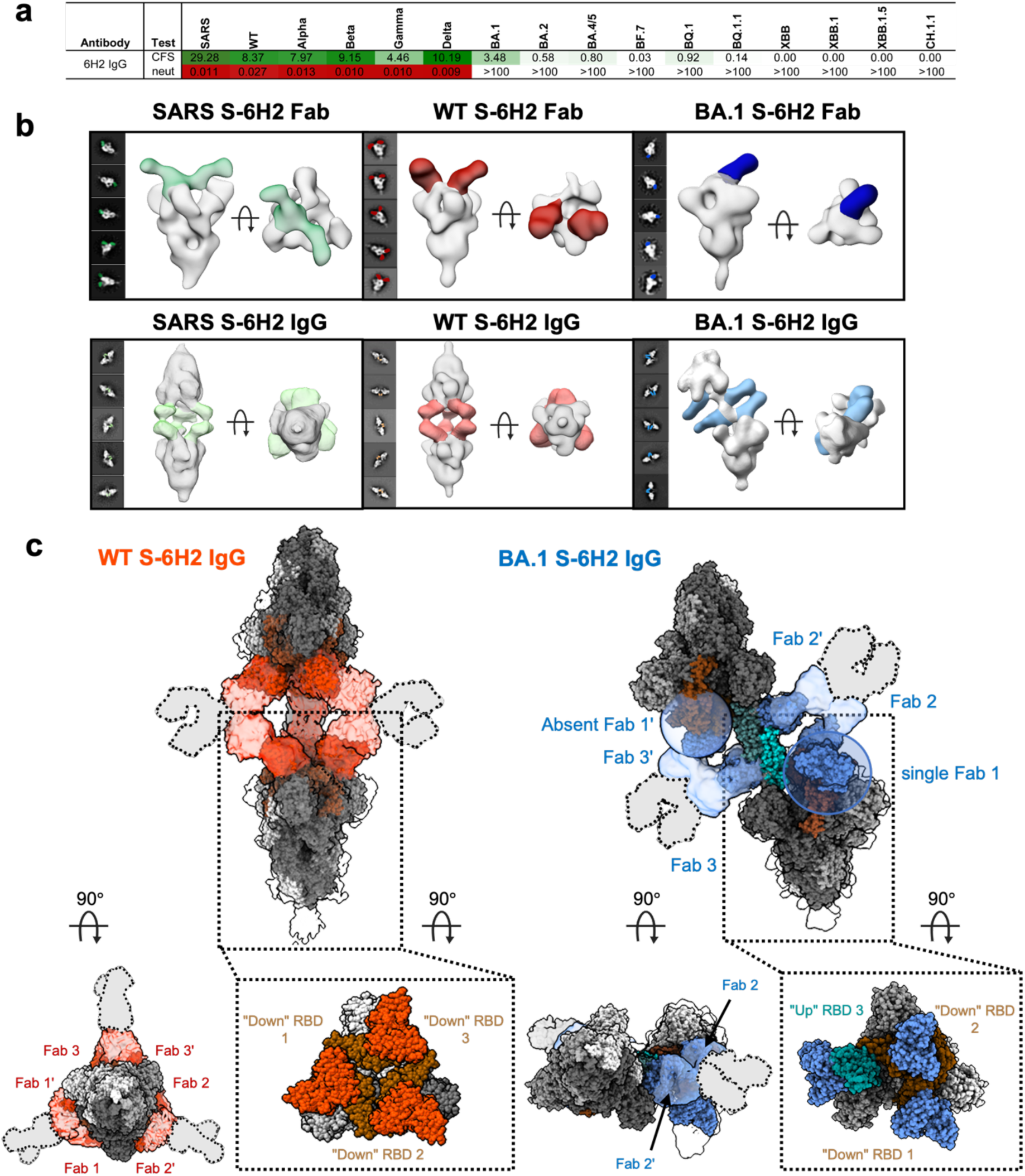
6H2 against SARS, WT and BA.1 variants with distinct dimer-trimer architectures. **a** The cross-binding ability and neutralizing potency (IC50) of 6H2 IgG to SARS and SARS-CoV-2 variants carried out by cell surface staining (CFS) and neutralization (neut) experiments. For cell surface staining, the diverse S of SARS and SARS-CoV-2 variants were expressed on the surface of HEK293T cells, stained with 6H2 IgG, and detected by flow cytometry. Total fluorescence intensity (TFI) of positively stained cells were calculated and normalized to represent the relative binding ability. For neutralization, the detection limit (the highest antibody concentration used for neutralization) was 100 μg/mL. Results of both experiments were derived from two independent repeats. **b** Representative two-dimensional (2D) classes and side and top views of composite figures from ns-EM analysis of 6H2 Fab or IgG complexed with S of SARS (green), WT (red) and BA.1 (blue) in a 3:1 molar ratio (Fab: S protomer). c The global cryo-EM structures of WT S-6H2 IgG and BA.1 S-6H2 IgG complexes. Two perpendicular views of complexes, with the Fab in Salmon (WT) or Cornflower blue (BA.1) and RBDs of S in Saddle brown, respectively. For BA.1 S-6H2 IgG complex, the region of Fab bound to the up non-bivalent down RBD domain showed weak density and were omitted in the final model (Absent Fab 1’).

To investigate the potential cause of the diminished neutralization of 6H2 IgG against Omicron variants from the structural perspective, we first employed ns-EM to characterize the global structure of 6H2 both in full-length IgG form and Fab form in complex with SARS, WT, Delta, and BA.1 S (Fig. 1b and Supplementary Table 1, 2). Upon binding with IgG, the complexes formed with SARS, WT, and Delta S adopted centered head-to-head dimer-trimer conformations (Fig. 1b and Supplementary Table 2). To our surprise, the conformations of IgG bound to Omicron BA.1 variant differed from that bound to WT, with 80% of the particles displaying non-symmetrical offset head-to-head dimer-trimer conformation, and only 20% forming the head-to-head dimer-trimer conformation (Fig. 1b and Supplementary Table 2). In our experimental conditions—ranging from a 3-fold (∼5.1 μM Fab to ∼1.7 μM S protomer) to 8-fold (∼13.6 μM Fab to ∼1.7 μM S protomer) excess of IgG and incubation times extending from 1 hour to 6h—we consistently recorded that nearly all S-6H2 IgG complex particles adopted this unique dimer-trimer configuration with 6H2 IgG (Supplementary Fig. 3 and 4). The reduction in binding percentage of 6H2 IgG from 94% (Fab/WT S protomer = 8:1) to 74% (Fab/BA.1 S protomer = 8:1) observed in the ns-EM analysis is consistent with the cell surface staining results (Fig. 1a). It is noted that the concentration used in the ns-EM experiments (0.3 mg/mL S protein) was significantly higher than the equilibrium dissociation constant (KD) value obtained from SPR (Supplementary Fig.5), however, given the inherent differences between these experimental approaches, complete complex formation was not observed with any of the samples^25, 26, 27^.

To further characterize the global binding modes of 6H2 in detail, we determined structures of the IgG bound to SARS, WT and BA.1 S using cryo-EM (Fig. 1c, Supplementary Fig. 6 and 7). While certain regions of cryo-EM structures appear blurry due to the intrinsic flexibility of IgG, 3D reconstructions clearly revealed two distinct dimer-trimer arrangements, consistent with the ns-EM results. For WT S-6H2 IgG, binding of 6H2 induced a centered head-to-head dimer-trimer structure where three IgGs cooperatively cross-linked two S trimers, with each IgG clamped two “down” RBDs from opposing S trimers, resulting in all RBDs being fully occupied (Fig. 1c). Upon binding to BA.1 S, 6H2 IgG induced a distinctive offset head to-head dimer-trimer structure, wherein two IgGs each cross-linked a “down” RBD (RBD-2 or 3’) and an “up” RBD (RBD-2’ or 3) from the opposite S, as shown in Fig. 1c. The third RBD of both S did not participate in inter-spike cross-linking, with one binding to a single Fab arm of 6H2 IgG and the other being unoccupied (Supplementary Fig. 7). Overall, the RBDs in each S exhibited one “up” and two “down” conformations, with a total of five 6H2 Fabs visible in the 3D reconstruction. Based on the distinct dimer-trimer structures, we propose that the complex, higher-order bivalent binding modes of 6H2 IgG could play an additional role in antibody neutralization. Furthermore, the observed loss of neutralization against Omicron variants could potentially be linked to these specific bivalent cross-linking modes of IgG.

### IgG structural adaptation accommodates diverse spatial arrangements

To quantitatively evaluate the changes in IgG angles and distances for the two distinct conformations, we employed a geometric analysis approach, establishing an auxiliary plane by connecting the same reference G104 residue in the light chain at the center of each Fab (Fig. 2a). In WT S-6H2 IgG structure, the auxiliary planes from two S trimers were parallel (∼0°) and the IgG approaching angle was nearly perpendicular (Fig. 2a). The distances between the Thr31 residues in the heavy chains of cross-linking Fabs were determined to be 102 Å, with the calculated angle being 85°, which suggested that the IgG was in a state close to an intact and relatively comfortable conformation. In BA.1 S-6H2 IgG, the same auxiliary plane revealed a larger angle of 32° between the surfaces, a non-perpendicular IgG approaching angle, and reduced Fab angle and distance (35° and 58Å, respectively) (Fig. 2b). This implies that, to bind to the RBDs on BA.1 S, 6H2 IgG must undergo twisting and deformation from its most stable state. Upon examining other potential inter-spike cross-linking modes involving two non-bivalent binding RBDs, we analyzed the distances and angles between Fabs and their docked regions. Our findings indicate that the current cross-linking mode is most favorable as the RBD1-RBD’ interaction resulted in either an obtuse angle or unsuitable distance (135 Å), which significantly deviates from the suggested inter-Fab distance of approximately 100 Å ^28^ (Fig. 2c). By aligning BA.1 S-6H2 IgG with ten published Omicron models (Supplementary Fig. 8), we observed flexibilities in the “up” RBD with the minimal and maximal changes labeled ① (Omi-native, pdb:7wvn)^29^ and ② (Fab: S3H3, pdb:7wk9)^29^ (Fig. 2d). In details, the “up” RBD in BA.1 S-6H2 exhibited a 10° rotation, while other views displayed 25° and 30° rotations. From the top view, RBDs in BA.1 S-6H2 IgG displayed a 70- and 80-degree angle with the native Omicron and Fab-bound structures, indicating that the RBD of BA.1 S-6H2 IgG appeared a larger open angle (Fig. 2d). Overall, 6H2 binding promoted the “up” RBD to overcome steric constraints and adopt a more “up” conformation.

**Fig. 2.**
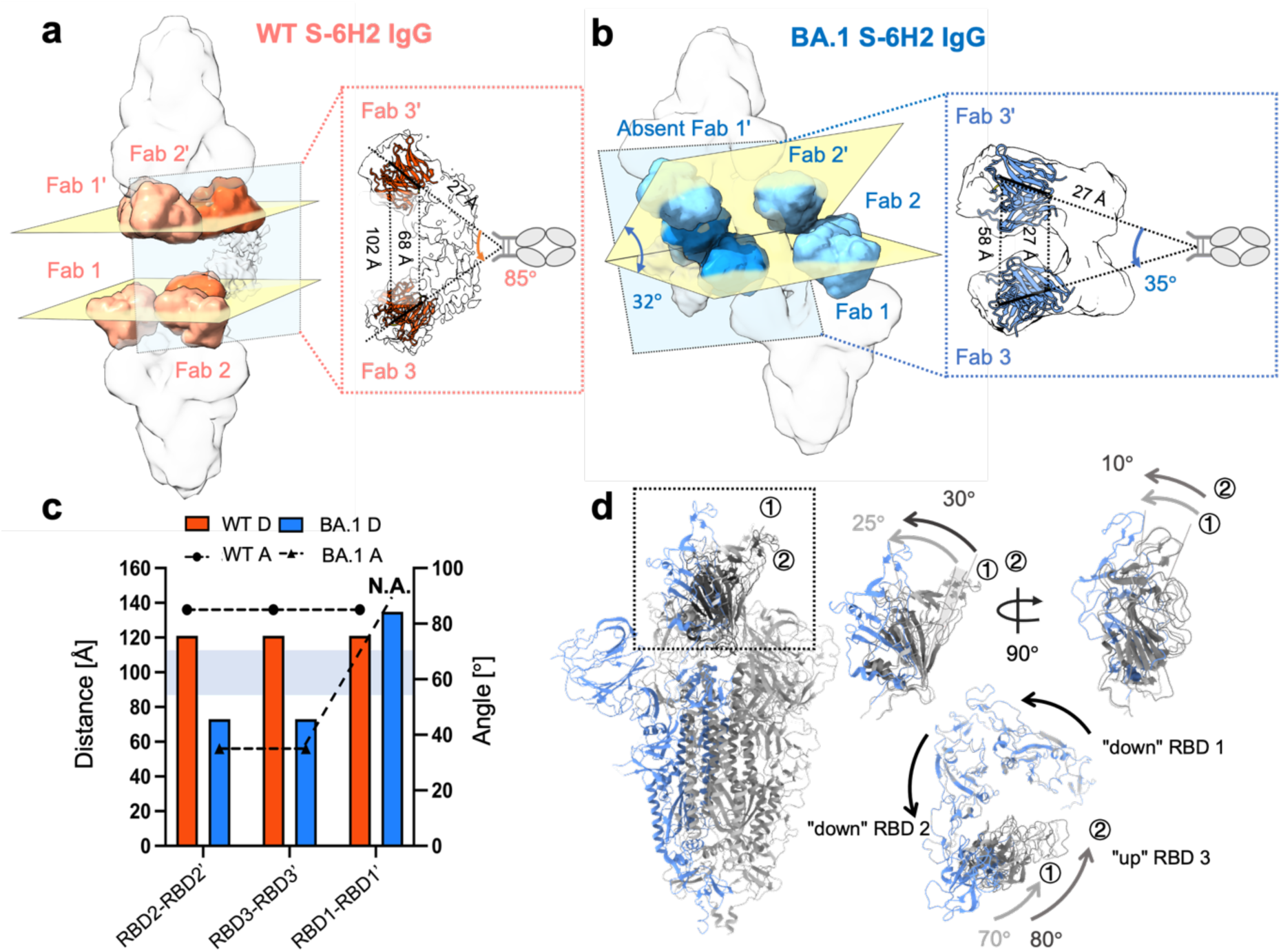
Spatial arrangements of two distinct dimer-trimer architectures. **a-b** Schematic illustration of the measured distance of two Fab domains, as well as the Fab-Fab angles from the two distinct dimer-trimer structures formed by WT (Red) and BA.1 (Blue) S. Two auxiliary planes (yellow) are defined by connecting the same reference G104 residue in the light chain at the center of each Fab and the measured angles between the two auxiliary planes are labeled. **c** The measured distances between the N343 residues of RBD domains, as well as the Fab-Fab angles of 6H2 IgG from distinct cross-linking mode formed by WT and BA.1. **d** The side view and top views of the overlaid protomers of BA.1 S-6H2 IgG, native BA.1-up RBD (① 7wvn) and BA.1-S3H3 (② 7wk9), indicating a larger open angle of RBD in BA.1 S-6H2 IgG.

### Binding mode is an underlying factor influencing the neutralization of IgG

To analyze the contact residues in detail, we built structures with local refined density maps that focused on the 6H2 Fab and “down” RBD region, at 3.55Å (SARS S-6H2 IgG), 3.81 Å (WT S-6H2 IgG) and 3.92 Å (BA.1 S-6H2 IgG) resolution (Supplementary Fig. 6 and Supplementary Table 3). Structural alignment of the RBD region was performed to identify commonalities and differences in epitopes among SARS S-6H2 IgG, WT S-6H2 IgG and BA.1 S-6H2 IgG complexes (Fig. 3a). The 6H2 binding was driven primarily by 11-residue-long complementarity-determining region 1 of the light chain (CDR-L1), 3-residue-long CDR-L2, 5-residue-long CDR-H1 and 16-residue-long CDR-H3 (Fig. 3a and Supplementary Fig. 9). As shown in Supplementary Fig. 9b, the loop 29-37 from CDR-L1, which occupied a large area of footprint, played a major role in 6H2 binding. Our analysis of the mutations that occurred in the epitope revealed that the presence of G446S within the epitope may be responsible for the large movement of loop 29-37 in BA.1 S-6H2 compared with the WT S-6H2. In contrast, although the N440K located near the CDR-L2 loop 55-57 in BA.1 S, it had minimal impact on the structures. Similarly, G339D also had little impact on the conformation of loop 99-104 in CDR-H3. In addition, the glycan at position N330 for SARS and N343 for SARS-CoV-2 variants played a crucial role in the targeting of 6H2 (Supplementary Fig. 9, 10 and 11). The glycan attached mostly with CDR-H3 (H100, V101, and L102) and exhibited a high degree of overlap among all models (Fig. 3a and Supplementary Fig. 9, 10, 11). Remodeling of electrostatic interactions at two mutation sites G339D and N440K may contribute to the dampened binding of 6H2 Fab with BA.1 S (Supplementary Fig. 12).

**Fig. 3.**
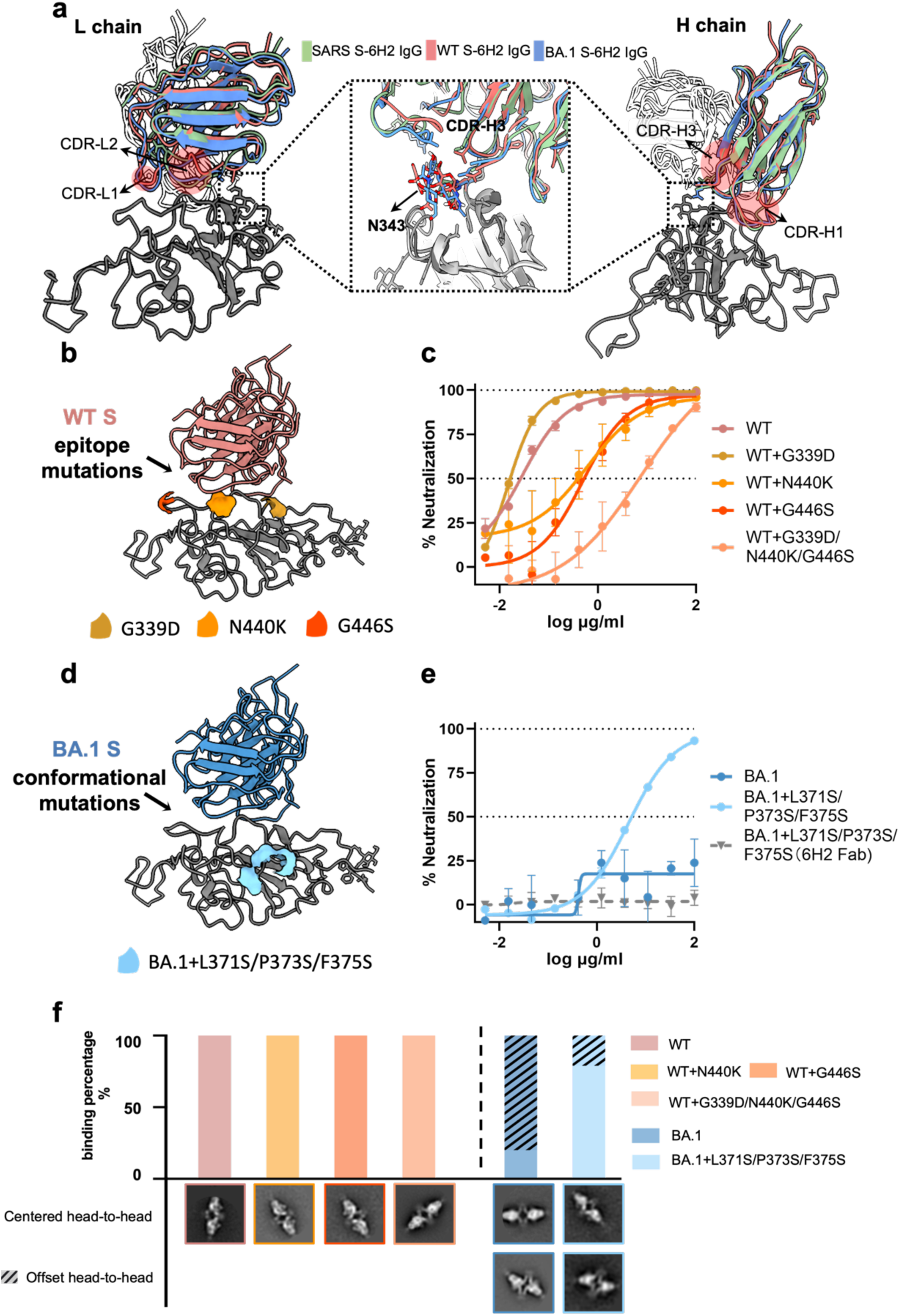
Distinct binding modes influence the neutralization of 6H2 IgG. **a** The same binding mode surrounded by N343 glycan among three distinct S-6H2 IgG complexes (green, red and blue, for SARS S-6H2 IgG, WT S- 6H2 IgG and BA.1 S-6H2 IgG). **b** Ribbon diagrams of WT S-6H2 IgG, with epitope mutations labelled and indicated by colored surface. **c** The neutralizing curve of 6H2 against WT, WT with G339D, N440K and G446S single-site mutations and G339D/N440K/G446S triple-site mutation with mean and s.e.m. labelled. **d** Ribbon diagrams of BA.1 S-6H2 IgG, with conformational mutations labelled and indicated by colored surface. **e** The neutralizing curve of 6H2 IgG and Fab against BA.1 and BA.1 with L371S/P373S/F375S triple-site mutation with mean and s.e.m. labelled. Results of both c and e were derived from two independent experiments, and each included two technical replicates **f** Ns-EM analysis denoting the proportion of centered or offset head-to-head dimer-trimers for each mutation (the total number refers to the number of complexes), and representative 2D class averages of distinct dimer-trimer structures shown in the bottom.

To discern the primary factors influencing antibody neutralization — namely, whether it is the epitope residues impacting the intrinsic affinity between the RBD and the 6H2 Fab, or the distinct cross-linking conformations that we previously identified — we designed two sets of mutations. 1) Epitope mutations including three single mutations (G339D, N440K and G446S) and G339D-N440K-G446S triple-site mutation on WT S (WT-M, Fig. 3b), and 2) conformational mutations encompass L371S-P373S-F375S sites on BA.1 S (BA.1-M, Fig. 3d), which are located distantly from the 6H2 epitope and have been reported to reduce the propensity for hydrophobic interactions and destabilize the one-RBD-up conformation of BA.1 S ^30^. The PsV neutralization assay revealed that epitope mutations on the WT-S led to a range of reduced neutralization potencies (IC50 values provided in Supplementary Fig. 13), yet they did not abolish neutralization entirely (Fig. 3c). Complete neutralization (∼100%) can be achieved at higher antibody concentrations (Supplementary Fig. 13). Notably, the conformational mutations L371S-P373S-F375S on BA.1 restored 90% neutralization at an antibody concentration of 100 µg/mL (Fig. 3e) and restored 100% neutralization at 1000 µg/mL (Supplementary Fig. 13).

Remarkably, ns-EM analysis of the BA.1-M S-6H2 IgG complex with conformational mutations showed a significant shift in the binding modes (Fig. 3f and Supplementary Fig. 14), with 79% centered head-to-head plus 21% offset, compared to a majority of 80% offset dimer in BA.1 S-6H2 IgG complex. This observation indicated that L371S-P373S-F375S triple mutations destabilized the one-RBD-up conformation of BA.1 S, leading to a centered head-to-head dimer-trimer structure with 6H2 IgG, where all the RBDs of the S-trimer adopt the “down” conformation. Meanwhile, the KD values of IgG decreased significantly from 1.71×10^-7^ M to 6.87×10^-9^ M after the transition from offset to centered head-to-head cross-linking, while KD values of Fab are comparable (2.28×10^-6^ M for BA.1 S and 3.80×10^-7^ M for BA.1-M) (Supplementary Fig. 5 and 15). Additionally, it was observed that the three single mutations as well as the G339D-N440K-G446S triple-site mutation on the WT epitope, consistently generated a centered head-to-head dimer-trimer structure with 6H2 IgG, directly attributed to a maximal neutralization efficacy of 100% (Fig. 3c and f). Crucially, we observed that while 6H2 IgG regained its neutralizing efficacy against BA.1-M, the 6H2 Fab remained non-neutralizing, indicating that, beyond the effect of the epitope, the bivalent binding of IgGs also plays a significant role in neutralization efficacy (Supplementary Fig. 13). Taken together, while epitope mutations undoubtedly influenced the affinity and neutralization potency of antibody, as expected, they are not the sole determinant. The conformational mutations introduced to BA.1 S profoundly altered the cross-linking modes. The resulting centered head-to-head configuration could lead to the restoration of neutralization efficacy.

### IgG binding modes can influence ACE2 accessibility

To gain further insight into the potential intrinsic mechanism underlying the binding mode-dependent variation in of 6H2 IgG, the binding behaviors between the preformed S-6H2 IgG complex to ACE2 were investigated by ns-EM, SPR, and FC assays. It is noted that the ACE2 construct used in this study included the Neck-domain of ECT region, which could mediate ACE2 dimerization as reported ^32, 33^ (Supplementary Fig. 16). When ACE2 was mixed with preformed WT S-6H2 IgG, the binding of ACE2 to preformed WT S-6H2 IgG was not observed in ns-EM 2D averages, and the head-to-head dimer structure of WT S-6H2 was preserved (Fig. 4a), demonstrating complete inhibition of ACE2 binding to WT S by 6H2 IgG. In contrast, when ACE2 was added to preformed BA.1 S-6H2 IgG, the plain offset dimer structure of BA.1 S-6H2 IgG could be hardly observed in ns-EM micrographs. While the majority formed larger single-particle complexes, some aggregation was observed in raw micrographs, potentially due to further cross-linking of BA.1 S-6H2 IgG by dimerized ACE2 monomer^34^ (Fig. 4b and Supplementary Fig. 16). Within the single-particle data, 2D classification revealed a significant decrease in the percentage of offset head-to-head dimers (4.9%) compared to the non-ACE2 group (51.2%), with a newly emerging particle class (38.3%) clearly demonstrating ACE2 binding to the offset head-to-head dimer-trimer (Supplementary Fig. 3 and 16). Ab-initio reconstruction was performed to identify the ACE2 binding site. The resulting 3D map revealed that two single ACE2 monomers simultaneously bound to the absent RBD 1’ and the monovalent site RBD 1 (Fig. 4c), while the other four RBDs retained the crosslinked by two IgGs (Fig. 4c and Supplementary Fig. 16). Overall, our findings demonstrated that the three IgGs fully crosslinked the six RBDs of WT S and tightly locked them in the “down” conformation, resulting in the formation of a stable symmetrical head-to-head dimer-trimer that effectively inhibited attachment or replacement of ACE2. In contrast, according to the cryo-EM analysis, BA.1 S-6H2 IgG complex displayed an asymmetrical dimer-trimer conformation where only four RBDs were crosslinked by IgG, exposing a single-bound RBD and an unoccupied RBD. As a result, the metastable offset dimer-trimer structure provided a pathway for ACE2 to approach, further inducing RBD “up” conformation and crowding out the monovalent binding IgG with lower binding affinity (Fig. 4c). We further determined the accessibility of ACE2 to preformed WT S-6H2 IgG and BA.1 S-6H2 IgG complexes with SPR (Fig. 4d-e). ACE2 was first immobilized on the chip, and the preformed complexes were injected. As shown in Figure 4d, responses after BA.1 S-6H2 IgG complexes injection were detected, compared to the null response after WT S-6H2 IgG complexes injection. The results revealed that 6H2 IgG effectively inhibited WT S binding to ACE2, whereas its impact on BA.1-S was little (Fig. 4e), suggesting that ACE2 was applicable to bind to the BA.1 offset dimer-trimer other than the WT head-to-head dimer-trimer. To better reflect the binding of ACE2 on the cell surface to S-6H2 complexes, we incubated the preformed WT S-6H2 IgG and BA.1 S-6H2 IgG complexes with HEK293T cells expressing ACE2, as shown in Fig. 4f. Using FC for quantification, we applied fluorescently labeled antibody that bound specifically to either the S protein or 6H2 IgG and compared the ACE2 availability of two different dimer structures on the cell surface (Fig. 4f-h and Supplementary Fig. 17). In particular, when 6H2 IgG was fluorescently labeled, only S-IgG complexes that were attached to ACE2 expressing cells contributed to FC signals, while the unoccupied S could not be stained, therefore did not affect our observations. When the ratio of 6H2 Fab to S protomer exceeded 1.25 in the mixture, the binding of WT S to the cell surface decreased markedly, accompanied by a decrease in the binding of 6H2 IgG. On the other hand, the binding of BA.1 S to the cell surface did not change with the ratio of Fab to S protomer, while the amount of cell surface-bound 6H2 IgG increased. As the proportion of antibody in the mixture increased, the amount of free S decreased while more S-6H2 IgG complexes formed. The decrease in antibody bound through WT S and increase in antibody bound through BA.1 S also confirmed that ACE2 on the cell surface can bind to the offset dimer-trimer of BA.1 S-6H2 IgG complexes.

**Fig. 4.**
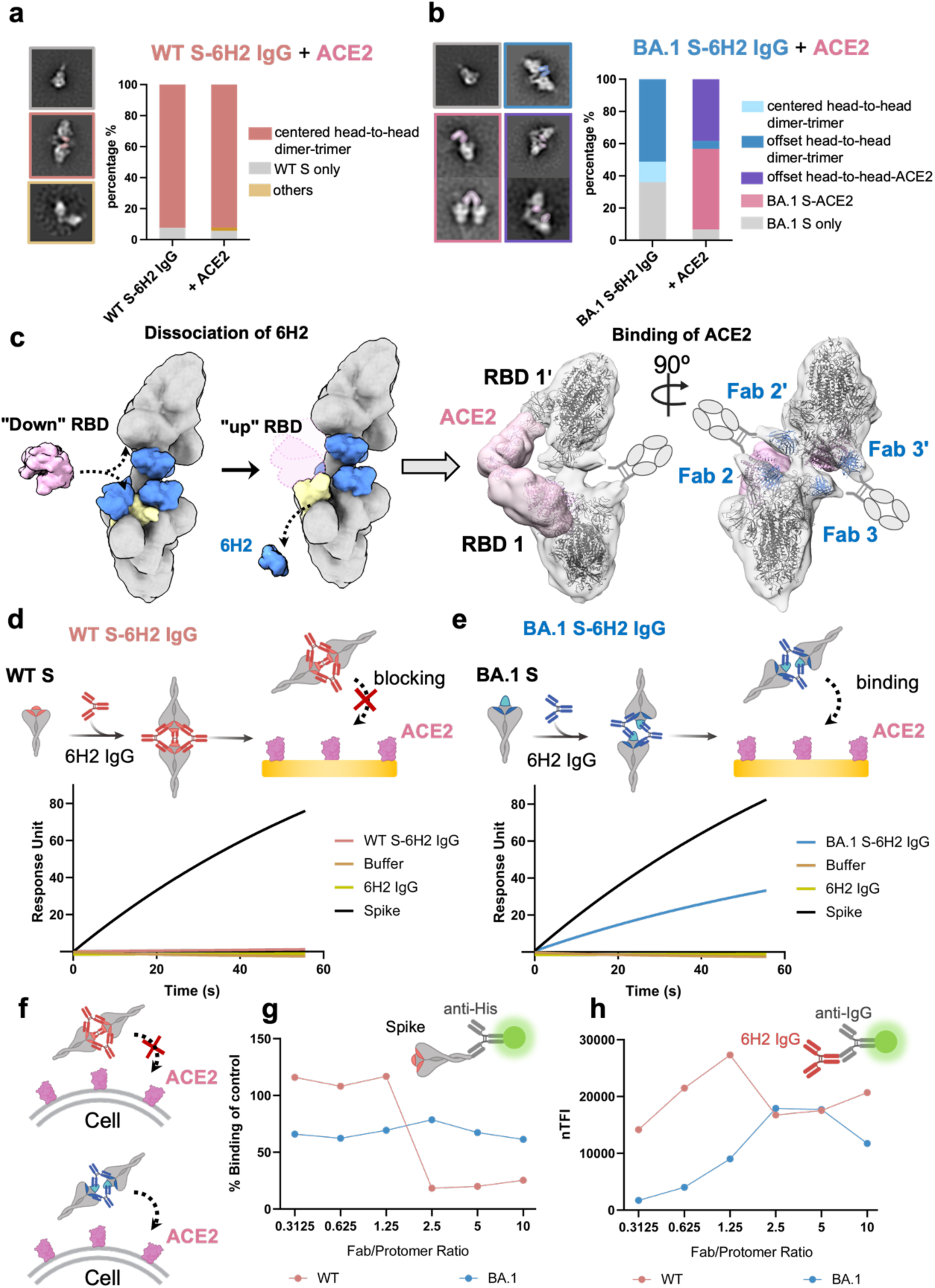
The accessibility of ACE2 to centered and offset head-to-head dimer-trimer architectures. **a-b** Ns- EM analysis denoting the proportion of each particle class formed by complexing WT S-6H2 IgG and BA.1 S- 6H2 IgG with ACE2, and representative 2D class averages shown in the left. **c** A flowchart displaying the binding process of ACE2 to BA.1 S-6H2 IgG. Fitting of the ACE2 (PDB:7sy0)^31^ and BA.1 S-6H2 IgG structure on the low-resolution 3D reconstruction of the complexes achieved from nsEM demonstrated the attachment of ACE2 to the BA.1 S-6H2 IgG. **d-e** SPR measurement of binding of WT S-6H2 IgG and BA.1 S-6H2 IgG complexes to ACE2. **f** Schematic of binding of WT S-6H2 IgG and BA.1 S-6H2 IgG to ACE2 on cell. **g-h** Total fluorescence intensity of percentage of S bound to 293T-ACE2 cells incubated with S-6H2 IgG, compared to the positive controls incubated with S alone, and the labeled 6H2 IgG attached to 293T-ACE2 cells through incubation of S- 6H2 IgG, were drawn into curves with results of different Fab/Protomer ratio, with (red) representing WT and (blue) representing BA.1. Results were derived from two independent experiments.

### Inter-spike binding attributes to the aggregation of viral particles

When extended to whole virus systems where multiple S present on the viral surface, the inter-spike cross-linking can potentially lead to the aggregation of viral particles^35^. To explore the impact of this specific mode of cross-linking on whole-virus neutralization, we designed a model system to mimic SARS-CoV-2 virus, employing 100 nm monodispersed Streptactin-coated polystyrene (PS) beads conjugated with SARS-CoV-2 S proteins (Fig. 5a). Our objective was to evaluate the 6H2 antibody at two distinct concentrations: 0.005 µg/mL, the minimal concentration previously utilized in our neutralization assays where neither WT nor BA.1 showed neutralization at this level, and 0.027 µg/mL, which corresponds to the IC50 value of 6H2 IgG against WT. We administered 6H2 IgG at these concentrations to beads conjugated with the WT and BA.1 S (WT-beads and BA.1-beads, respectively), and characterized the degree of aggregation. As shown in Fig. 5b and Supplementary Fig. 18, ns-EM micrographs showed no evidence of aggregation at the lower concentration of 0.005 µg/mL for both WT and BA.1-beads. However, at the elevated concentration of 0.027 µg/mL, a significant aggregation was observed in the WT-beads-6H2 IgG complexes. In comparison, the BA.1-beads-6H2 IgG complex displayed minimal aggregation, with additional ns-EM raw micrographs provided in Supplementary Fig. 18. To our pleasant surprise, the BA.1-M with triple conformational mutations L371S/P373S/F375S exhibited aggregation patterns similar to WT. This observation further supported our hypothesis that the highly stable centered cross-linking mode contributed to a greater degree of aggregation, ultimately facilitating the restoration of neutralization.

**Fig. 5.**
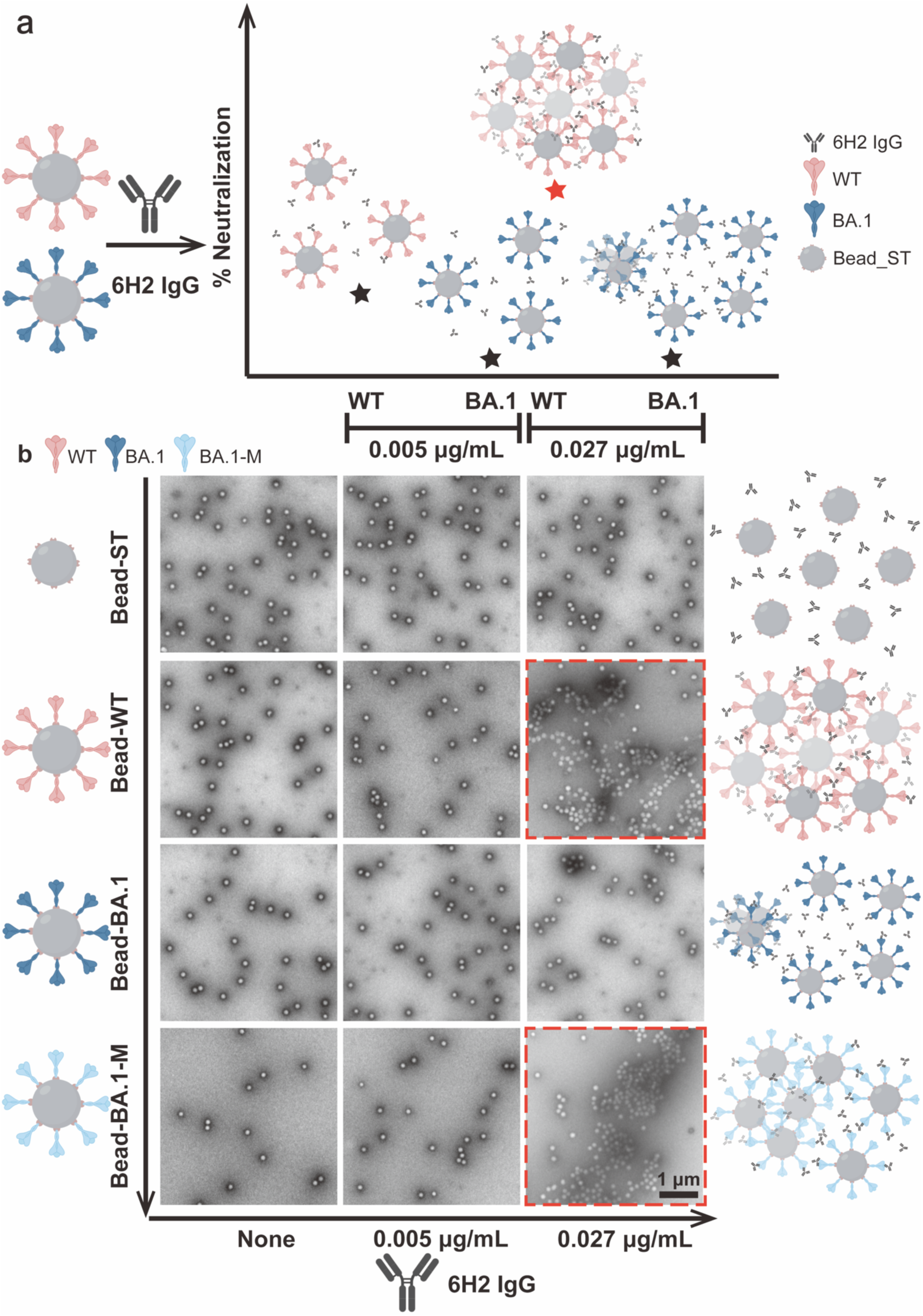
The aggregation of S protein modified beads induced by inter-spike cross-linking of 6H2 IgG. **a** Schematic illustration of beads modified with S protein upon addition of 6H2 resulting in aggregation. **b** Representative ns-EM raw micrographs characterizing the extent of aggregation in beads modified with WT, BA.1 S and BA.1-M S after the addition of 6H2 IgG. Additional representative images of this experiment are also shown in the Supplementary Fig. 18. All results were repeated at least twice in independent experiments. Scale bar: 1 μm. The scale bar in the last image is applicable to all other images in the same panel.

This 100 nm whole virion model system probes the collective behavior of multiple S proteins. It is recognized that not every S on the spherical virion surface is capable of forming head-to-head dimers. This, in particular, highlights the critical role of stability in these complexes, serving as anchoring points for viral aggregation. When adopting a centered binding mode, full stoichiometry binding facilitates maximal synergy, culminating in the formation of the most stable complexes. Conversely, the offset cross-linking mode, with sub-stoichiometry, tends to yield significantly fewer stable complexes (Fig. 1c and 5b). This difference in synergy directly influences the degree of viral aggregation, and ultimately impacts the accessibility of viruses to cellular receptors. With the same concentration of 6H2 present, the less stable offset head-to-head complexes formed between BA.1 S and 6H2 IgG led to reduced aggregation. These observations altogether underscore the intricate relationship between cross-linking architecture, aggregation, and 6H2 antibody immune evasion mechanisms.

## DISCUSSION

The multivalent interactions play a vital role in a wide range of biological recognition and signaling processes, including immune recognition, cell adhesion, virus-host interactions, and protein-protein interactions^35^. In particular, the antibody-antigen interaction is a classic example of multivalent interactions^36^. Herein, we have observed two unique dimer-trimer architectures using a bivalent W328-6H2 IgG in complex with SARS, WT, and BA.1 S. In our observation, regardless of the excess fold of IgG and incubation time, the crosslinked dimer structure always preferentially forms instead of each S independently binding three IgG molecules. Such inter-spike crosslinking behaviors have also been previously observed with other antibodies complexed with S^8, 12, 24, 37, 38^, suggesting that the propensity for bivalent IgG to mediate the formation of crosslinked dimer structures is a common feature of antibody-antigen interactions. Furthermore, summing up our observations from ns-EM, SPR, cell surface staining, and neutralization experiments, it is evident that such dimer-trimer architectures exhibit significantly enhanced binding avidity and neutralization of bivalent 6H2 IgG compared to its monovalent Fab. This improvement can be attributed to the synergistic enhancement effects of multivalent interactions, which can be understood from two perspectives^39^. Thermodynamically, the simultaneous binding of multiple interactions leads to a more significant change in free energy (Δ*G*), resulting in superior binding avidity of bivalent IgG (according to Δ*G* = −*k_B_T* ln *K*, where *K* is the binding constant). From a kinetic view, while the diffusion-controlled association rate constant ( *k*_a_) of IgG is similar to that of monovalent Fab, the dissociation rate constant (*k*_d_) of IgG is significantly decreased due to the higher activation energy of dissociation, also contributing to the enhanced binding avidity of 6H2 IgG^40, 41, 42, 43^.

Neutralization is widely defined as the loss of infectivity which ensues when antibody molecule(s) bind to a virus particle. Antibody-mediated neutralization encompasses a broad spectrum of mechanisms^23^, primarily through preventing the attachment of viral particles to target cells by blocking the binding of host cell receptors^9^, inhibiting conformational changes of the S protein^44, 45^, inducing disassembly of the S protein^23^ and facilitating aggregation of viral particles^35^. In the case of 6H2 antibody, the formation of viral aggregation through inter-spike cross-linking could lower the probability of viral infection to host cells^46^ by trapping large portions of surface S protein within aggregates, block cellular attachment, and even impede subsequent endosomal fusion and egress — as similarly observed with influenza antibody HC19^47^, Chikungunya virus (CHIKV) antibody CHK-124^35^, human immunodeficiency virus (HIV) antibody 2F5^48^ and Rhinovirus (RV) antibody MAb 15^49^. Conversely, the significantly reduced degree of aggregation of BA.1 and 6H2 leads to a higher count of non-aggregated effective viral entities that remain unbound or form asymmetric offset dimer structures with 6H2. The asymmetric spatial arrangement of BA.1 S no longer supports the bivalent crosslinking of all three IgGs as observed in the centered dimer structures. As a result, the reduced occupancy of RBDs facilitates ACE2 approaching and potentially aids the immune escape of Omicron variants.

In summary, by utilizing full-length IgG—the biologically relevant form of antibodies that offer protection following infection or vaccination—our study has bridged a gap in understanding multivalent binding modes in spike-antibody cross-linking interactions. This research underlines that beyond the binding affinity between the RBD epitope and Fab, the global binding mode of IgG plays an additional role in influencing the antibody’s avidity and neutralization capabilities. These insights should be integral to the considerations for designing novel therapeutic antibodies and vaccines.

## Methods

### Cell lines

HEK293T cells (ATCC, CRL-3216) and 293T-hACE2 cells were maintained at 37 °C in 5% CO_2_ in Dulbecco’s minimal essential medium (DMEM) containing 10% (v/v) heat-inactivated fetal bovine serum (FBS) and 100 U/mL of penicillin–streptomycin. 293T-hACE2 cells were constructed to stably express human ACE2 and susceptible to SARS-CoV-1 and SARS-CoV- 2 infection. FreeStyle 293F cells (Thermo Fisher Scientific, R79007) were maintained at 37 °C in 5% CO_2_ in SMM 293-TII expression medium (Sino Biological, M293TII).

### Antibody and Fab fragment production

6H2 is a potent neutralizing mAb initially isolated from SARS infected patients by our group^24^. All the IgG heavy and light chain variable regions were cloned into expression vectors encoding full length constant region to produce IgG1 antibodies as previously described^50^. Briefly, genes encoding the heavy and light chains of antibodies were transiently transfected into HEK 293 F cells using polyethylenimine (PEI) (Sigma). After 96 h, antibodies in the supernatant were collected and captured by Magnetic Protein A-Sepharose (GE Healthcare). Bound antibodies were eluted by solution buffer Glycine pH 2.0 and further purified by gel filtration chromatography using a Superdex 200 High Performance column (GE Healthcare). To produce Fab fragments, antibodies were cleaved using Protease Lys-C (Roche) with an IgG to Lys-C ratio of 4000:1 (w/w) in 10 mM EDTA, 100 mM Tris-HCl, pH 8.5 at 37 °C for approximately 12 h. Fc fragments were removed using Protein A Sepharose. Antibody and Fab concentration were determined by Nanodrop 2000 Spectrophotometer (Thermo Scientific).

### Protein expression and purification

The human codon-optimized gene of recombinant trimeric spike of SARS-CoV-1 spike-His-Strep-FLAG (residues 1-1195, K968P, V969P) and Delta spike-His-Strep-FLAG (residues 1- 1208, GSAS, K986P and V987P) were synthesized in GeneScript and cloned into pCAGGS vector, and SARS-CoV-2 spike-His-Strep-II (residues 1-1208, GSAS, F817P, A892P, A899P, A942P, K986P and V987P) and BA.1 variant spike-His-Strep-II (residues 1-1208, GSAS, F817P, A892P, A899P, A942P, K986P, V987P) were synthesized in Tsingke Co., LTD and cloned into SARS-CoV-2 S HexaPro (addgene: 154754). The single and triple-site mutations were introduced into the recombinant SARS-CoV-2 wildtype and BA.1 spike vectors constructed above. Gene expressing recombinant human ACE2 peptidase domain with flag and strep tags (residues S19 to S742) was cloned into pVRC vector.

All recombinant proteins were produced by transfecting the expression vectors into the FreeStyle 293F cells. Ninety-six hours after transfection, trimeric spikes and ACE2 secreted into the cell supernatant were harvested and captured by Strep-Tactin Sepharose and further purified by gel-filtration chromatography using Superdex 200 High-Performance columns (GE Healthcare). The final protein concentrations were determined using a nanodrop 2000 Spectrophotometer (Thermo Scientific).

### Production of pseudoviruses

Sequences of spike that SARS-CoV-1 (GenBank: GCA_000864885.1) and SARS-CoV-2 (GenBank: MN908947.3) pseudoviruses used were the prototype strains. SARS-CoV-2 variants were prepared as follows. The Alpha variant (Pango lineage B.1.1.7, GISAID: EPI_ISL_601443) included a total of 9 reported mutations in the spike protein: 69-70del, 144del, N501Y, A570D, D614G, P681H, T716I, S982A and D1118H. The Beta variant (Pango lineage B.1.351, GISAID: EPI_ISL_700450) included 10 identified mutations in the spike, namely, L18F, D80A, D215G, 242-244del, S305T, K417N, E484K, N501Y, D614G and A701V. The Gamma variant (Pango lineage P.1, GISAID: EPI_ISL_792681) had 12 reported mutations in the spike comprising L18F, T20N, P26S, D138Y, R190S, K417T, E484K, N501Y, D614G, H655Y, T1027I and V1176F. The Delta variant (Pango lineage B.1.617.2, GISAID: EPI_ISL_1534938) included 10 reported mutations in the spike, i.e. T19R, G142D, 156-157del, R158G, A222V, L452R, T478K, D614G, P681R, D950N. The BA.1 variant (Pango lineage BA.1, GISAID: EPI_ISL_6752027) was constructed with 32 mutations in the spike comprising A67V, Δ69-70, T95I, G142D/Δ143-145, Δ211/L212I, ins214EPE, G339D, S371L, S373P, S375F, K417N, N440K, G446S, S477N, T478K, E484A, Q493R, G496S, Q498R, N501Y, Y505H, T547K, D614G, H655Y, N679K, P681H, N764K, D796Y, N856K, Q954H, N969K, and L981F. The BA.2 variant (Pango lineage BA.2, GISAID: EPI_ISL_8515362) was constructed with 29 mutations in the spike as follows: T19I, 24-26del, A27S, G142D, V213G, G339D, S371F, S373P, S375F, T376A, D405N, R408S, K417N, N440K, S477N, T478K, E484A, Q493R, Q498R, N501Y, Y505H, D614G, H655Y, N679K, P681H, N764K, D796Y, N969K and Q954H. BA.4 and BA.5 variants (Pango lineage BA.4, GISAID: EPI_ISL_12559461) shared the same sequence on spike, and the variant was constructed based on BA.2 variant with additional of Δ69-70, L452R and F486V substitutions and the removal of the Q493R substitution. BF.7 (Pango lineage BF.7, GISAID: EPI_ISL_15429967), BQ.1 (Pango lineage BQ.1, GISAID: EPI_ISL_15458271), BQ.1.1 (Pango lineage BQ.1.1, GISAID: EPI_ISL_15458263) variants were then constructed based on BA.4 variant with additional of R346T, or K444T and N460K, or all of the three mutation sites. XBB variant (Pango lineage XBB, GISAID: EPI_ISL_15601178) contained 13 more mutation sites of V83A, Δ145, H146Q, Q183E, V213E, G339H, R346T, L368I, V445P, G446S, N460K, F486S, F490S, and the removal of Q493R, compared to BA.2 variant. G252V was added to XBB, which formed XBB.1 (PANGO lineage XBB.1, GISAID: EPI_ISL_15596825). F486S of XBB.1 was substituted by F486P to form XBB.1.5 (PANGO lineage XBB.1.5, GISAID: EPI_ISL_16283160). CH.1.1 (Pango lineage CH.1.1, GISAID: EPI_ISL_16702429) was constructed based on the BA.2 sequence, with addition mutations of: K147E, W152R, F157L, I210V, G257S, G339H, R346T, K444T, G446S, L452R, N460K, and F486S, and a removal of Q493R. The single and triple-site mutations were introduced into the pcDNA3.1 vector encoding SARS-CoV-2 wildtype or BA.1 Spike using QuickChange site-directed mutagenesis (Agilent 210519) according to the instructions. The full-length genes of spike variants were synthesized by Genwiz, Inc. and verified by sequencing. Pseudoviruses were generated by co-transfecting HEK-293T cells (ATCC) with human immunodeficiency virus backbones expressing firefly luciferase (pNL4-3-R-E-luciferase) and pcDNA3.1 vectors encoding either wildtype or variant spike proteins. Viral supernatant was collected 48 to 72 h after transfection, centrifuged to remove cell debris, and stored at −80°C until use.

### Antibody neutralization using pseudoviruses

Antibodies were diluted before being mixed with the pseudoviruses at 37°C for 1 h and added onto HeLa-hACE2 cells. After 48 h, the infected cells were lysed, and luciferase-activity measured. The percentage of neutralization was determined in comparison to virus control. Antibodies were diluted starting from 10 μg/mL for W328-6H2 IgG against SARS and WT, Alpha, Beta, Gamma, Delta pseudoviruses and from 100 μg/mL for the other cases. Half-maximal inhibitory concentration (IC50) of mAb was calculated by fitting the four-parameter dose inhibition response model with Graphpad Prism 8.0.

### Binding of SARS antibodies to cell surface-expressed spike glycoproteins

The entire procedure was conducted as previously published^51^. Specifically, HEK293T cells were transfected with expression plasmids encoding the various full-length spikes at 37 °C for 36 h. Cells were digested from the plate with trypsin and distributed into 96-well plates. Cells were washed twice with 200 µL staining buffer (PBS with 2% heated-inactivated fetal bovine serum) between each of the following steps. First, cells were stained with test antibody of IgG or Fab form of 1 μg/mL for SARS-CoV-2 variants’ and single/triple-site mutants’ cross-binding capacity detection, or serial dilution starting from 50 μg/mL for WT, WT with N440K and G446S single-site mutant, and BA.1 variant binding capacity comparison. A control S2- specific mAb (MP Biomedicals, Singapore 08720401) was used as positive control. The first step was carried out at 4 °C for 30 min in 100 μL staining buffer. Then, Alexa FluorTM 647- labeled anti-human IgG (H+L) (Invitrogen A-21445), or Alexa FluorTM 647-labeled anti-mouse IgG (H+L) (Invitrogen A-21236) was added in 50 μL staining buffer at 4 °C for 30 min. After extensive washes, the cells were resuspended and analyzed on a BD LSRFortessa (BD Biosciences, USA) using FlowJo 10 software (FlowJo, USA). Mock-transfected HEK293T cells were stained as negative control. Normalized total fluorescence intensity (nTFI) values were calculated by multiplying the number of positive cells in the selected gates with their mean fluorescence intensity, relative to that of S2 specific antibody.

### Antibody binding kinetics measured by SPR

The binding kinetics of IgG or Fab form of 6H2 to SARS-CoV-1, SARS-CoV-2 mutated, mutated or variant spikes were analyzed by SPR (Biacore 8K, GE Healthcare). Specifically, recombinant spikes were covalently immobilized to a CM5 sensor chip via amine groups in 10 mM sodium acetate buffer (pH 4.0) for a final response unit around 1000. Serial dilutions of W328-6H2 in IgG or Fab form flowed through the sensor chip system. The resulting data were fitted to a 1:1 binding model using Biacore Evaluation Software (GE Healthcare).

### ACE2 accessibility of spike-6H2 complex detected by SPR

Recombinant ACE2 was covalently immobilized to a CM5 sensor chip via amine groups in 10 mM sodium acetate buffer (pH 4.0) for a final response unit of 5000. Mixtures of spikes and IgG or Fab form of 6H2 flowed through the sensor chip system. Responses of spikes or antibody alone were also recorded as control.

### Binding of W328-6H2-spike mixture on the cell surface measured by flow cytometry

W328-6H2 IgG and 0.6 μM WT/BA.1 spike were incubated with a serial Fab/Protomer ratio from 10 to 0.3125 for 4 hours. HEK293T-ACE2 cells were digested from the plate with trypsin and distributed into 96-well plates. Cells were washed twice with 200 µL staining buffer (PBS with 2% heated-inactivated fetal bovine serum) between each of the following steps. Firstly, W328-6H2-spike complex mixtures were 30-fold diluted and incubated with HEK293T-ACE2 cells in 100 μL straining buffer at 4 °C for 45 min. Then, Alexa FluorTM 647-labeled anti-human IgG (H+L) (Invitrogen A-21445) or Streptavidin PE (12431787) was each added in 50 μL staining buffer at 4 °C for 30 min to label the antibody or spike. After extensive washes, cells were resuspended and analyzed with BD LSRFortessa (BD Biosciences, USA) using FlowJo 10 software (FlowJo, USA). Cells incubated with spike alone were treated as positive controls for the spike labelled samples, while cells incubated with antibody alone were treated as negative controls. Normalized total fluorescence intensity (nTFI) of spike or antibody was acquired by multiplying the percentage of positive cells with the mean fluorescence intensity. Percentage of spike binding to the cell surface were calculated by the following formula: (nTFI of spike labelled sample) / (nTFI of spike labelled positive control) × 100%.

### Streptactin proteins modification onto polystyrene (PS) beads

Streptactin (ST) proteins were covalently conjugated with 100 nm carboxy-functionalized PS beads through an EDC/NHS condensation reaction. An EDC/Sulfo-NHS solution (EDC: 100 mM, Sulfo-NHS: 80 mM) was prepared in a MEST buffer solution (100 mM MES + 0.05% Tween 20, pH 5.89). Next, 20 μL of carboxy-functionalized PS beads suspension was added to 200 μL of EDC/Sulfo-NHS solution and incubated for 20 minutes at room temperature (RT) to activate the carboxy groups on the surface of the PS beads. The resulting solution was then washed with PBST (PBS + 0.05% Tween 20, pH 7.43) to eliminate any unreacted EDC, Sulfo-NHS, and byproducts. Then, ST proteins were introduced to bind them covalently to the beads. The mixture underwent incubation at 4 ℃ with vortexing for 4 hours, and the reaction was quenched by adding Tris-HCl. Finally, transmission electron microscopy (TEM, HITACHI HT-7700) was employed to ascertain the monodisperse of the beads.

### Bead_based aggregation assay

Each mimic SARS-CoV-2 consists of a single Streptactin-coated PS bead and SARS-CoV-2 S bearing the Twin Strep tag. The ST-coated beads and either the WT, BA.1, or BA.1-M S are blended and incubated at RT for 4 hours without any further washing steps. Due to the high binding affinity between Streptactin and Twin strep tag, the spike proteins are conjugated onto the surface of ST-functionalized PS beads with a nominal size of 100 nm (similar to the size of SARS-CoV-2 virus). Subsequently, 6H2 IgG at varying concentrations (0.005 μg/mL, 0.027 μg/mL) was introduced into the WT/BA.1/BA.1-M-beads solution and incubated at RT for 1 hour. Finally, the beads suspensions were diluted with TBS and deposited on a carbon-coated electron microscope grid (200 with mesh copper, Zhongjingkeyi Films Technology Co.,Ltd), followed by negatively staining with 2% uranyl acetate. The distribution states of beads were performed on transmission electron microscope at 80 KV.

### Negative-stain electron microscopy

Grid preparation, image processing, and raw data analysis followed a similar protocol described previously^24^. Briefly, S proteins were complexed with IgG/Fab at 3× molar excess of Fab/protomer ratio and incubated at room temperature 1 h or 6 h. Due to the low binding percentage, crosslinker was added to BA.1 S-6H2 Fab to reconstruct 3D conformation (Supplementary Fig. 19). For WT S-6H2 IgG-ACE2 or BA.1 S-6H2 IgG-ACE2, S proteins were complexed with 6H2 IgG at 3× molar excess of Fab/protomer ratio and incubated at room temperature for 5 hours. Then ACE2 was added at 3× molar excess of ACE2/protomer ratio and incubated at room temperature 1h or 6h. The final concentration for complexes in incubation solution was ∼0.30 µg/µL. Complexes were diluted to 0.015 µg/mL in 1× Tris-buffered saline and 3.5 µL applied to a 200 mesh Cu grid (Zhongjingkeyi Films Technology Co.,Ltd) blotted with filter paper, and stained with 0.7% uranyl acetate (Electron Microscopy Sciences. 22400. USA). Micrographs were collected on a JEOL JEM-2100F high resolution transmission Electron microscope operating at 200 kV with a camera at ×30,000 magnification using the serial EM automated image collection software. EM-map reconstruction was performed using relion 3.1.3^52^, and the maps were aligned and displayed using Chimera. The ns-EM maps are available in the EMD database (www.emdataresource.org) and access numbers are listed in Table S2 (will update after EMDB submission).

### Cryo-EM sample preparation and data collection

Cryo-EM single-particle analysis was carried out following the detailed protocols published previously^53, 54^. Briefly, Purified SARS, WT and BA.1 S were mixed with 6H2 IgG at 3× molar excess of S protomer ratio and incubated at room temperature overnight. The final concentrations of the mixtures were 0.4 mg/mL in TBS buffer. Prior to cryo-EM grid preparation, the EM grids (Quantifoil grid, Cu 300 mesh, R2/1) were glow-discharged for 20 s using a plasma cleaner. Then, 3.5 μL complexes were applied to the grids and the grids were then blotted for 2 seconds with filter paper in 100% relative humidity and 22 ℃ and plunged into the liquid ethane to freeze samples using FEI Vitrobot system (FEI).

Cryo-EM data were collected using FEI Titan Krios (Thermo Fisher Scientific) electron microscope operating at 300 kV with a Gatan K3 Summit direct electron detector (Gatan Inc.). Automated data collection was carried out at a magnification of 29,000 with a pixel size of 0.97 Å and at a defocus range between 1.2-1.5 μm using SerialEM software^55^. Each movie had a total accumulate exposure of 50 e^-^/Å^2^ fractionated in 32 frames.

### Cryo-EM image processing

All the data processing was carried out using either modules on, or through, cryoSPARC v.3.3.1^56^. Parameters of contrast transfer function (CTF) were estimated by using Gctf11. All micrographs then were manually selected for further particle picking upon ice condition, defocus range and estimated resolution.

### Model building and refinement

Initial models were generated by fitting spike coordinates from Protein Data Bank (PDB) 7w9k (for SARS S-6H2 IgG), 7wvo (for WT S-6H2 IgG) and 7wv1 (for BA.1 S-6H2 IgG) into the cryo-EM maps using University of California San Francisco Chimera^57^. N-linked glycans were hand-built into the density where visible, and Several rounds of iterative manual and automated model building and relaxed refinement were performed using Coot 0.9.4^58^ and Rosetta^59^. Models were validated using EMRinger^60^ and MolProbity^61^ as part of the Phenix software suite^62^. Heavy and light chain sequences were exported into a Fasta file and blasted on IMGT described previously^24^. Final refinement statistics and PDB deposition codes for generated models can be found in SI Appendix, Table S3 (will update after EMDB and PDB submission). And the figures were generated using UCSF Chimera and ChimeraX.

## Data availability

Atomic models and EM densities have been deposited into the PDB and EMDB as follows: EMD-36257(Immune complex of W328-6H2 IgG binding the RBD of SARS-CoV-1 2p spike protein), EMD-35961(Immune complex of W328-6H2 Fab binding the RBD of SARS-CoV-2 WT 6p spike protein), EMD-36267(Immune complex of W328-6H2 IgG binding the RBD of SARS-CoV-2 WT 6p spike protein), EMD-35962(Immune complex of W328-6H2 Fab binding the RBD of Omicron BA.1 6p spike protein added BS3 crosslinker), EMD-35963(Immune complex of W328-6H2 IgG binding the RBD of Omicron BA.1 6p spike protein), EMD- 35964(Immune complex of W328-6H2 Fab binding the RBD of Omicron BA.4/5 6p spike protein), EMD-35969(Immune complex of W328-6H2 IgG binding the RBD of Omicron BA.4/5 6p spike protein), EMD-35970(Human ACE2 binding the complex of Omicron BA.1 6p spike protein and W328-6H2 IgG), EMD-35986(Cryo-EM structure of SARS-CoV-1 2p spike protein in complex with W328-6H2 IgG), EMD-35995(Cryo-EM structure of SARS-CoV-2 WT 6p spike protein in complex with W328-6H2 IgG), EMD-36058(Cryo-EM structure of Omicron BA.1 6p spike protein in complex with W328-6H2 IgG), 8JAG/EMD- 36113(Locally refined Cryo-EM structure of SARS-CoV-1 2p RBD in complex with W328- 6H2), 8JAP/EMD-36121(Locally refined Cryo-EM structure of SARS-CoV-2 WT RBD in complex with W328-6H2), 8JAM/EMD-36122(Locally refined Cryo-EM structure of Omicron BA.1 RBD in complex with W328-6H2). Ns-EM structures have also been provided in Supplementary Table 2. All data generated or analyzed during this study are available within the paper and the Supplementary Information files.

## Acknowledgements

The authors would like to thank the National Key R&D Program of China (2022YFA1206400), the National Natural Science Foundation of China (22277017, 92169205, 82150205, and 32270983), the National Key Plan for Scientific Research and Development of China (2021YFC0864500 and 2022YFC2604103, 2022YFC2303403), the Wanke Scientific Research Program (20221080056), Tencent Foundation, Shuidi Foundation, and TH Capital for financial support. N.L. was supported by CAMS Innovation Fund for Medical Sciences (CIFMS 2019-I2M-5-018).

## Author contributions

Y.Y., L.Z. and T.L. conceived and designed the study. X.N., Y.L., R.Z., R.W. and N.L. performed most of the experiments, with assistance from Y.W., B.Z., Y.C. and Z.W. X.N. solved and analyzed the ns-EM and cryo-EM structures of the antibody-spike complex, with assistance from Y.L., B.Z., J.L. and J.Z. Y.W., Z.W. and N.L. provided assistance with production of antibody and trimeric spike protein. N.L. performed the bead_based aggregation assay. W.C., Q.Z., X.S., C.Z., C.C., Z.L., Y.Z., D.L. and X.W. provided additional technical or intellectual assistance. X.N., Y.L., R.Z., R.W., N.L., T.L., L.Z. and Y.Y. had full access to data in the study, generated figures and tables, and take responsibility for the integrity and accuracy of the data presentation. Y.Y., X.N. and Y.L. wrote the manuscript. All authors reviewed and approved the final version of the manuscript.

