## Supplementary Materials for "Distinct Binding Modes of Inter-Spike Cross-Linking Suggest a Supplementary Mechanism for SARS-CoV-2 Antibody Neutralization"

### Contents


| | |
|---|---|
| Supplementary Fig. 1 | 3 |
| Supplementary Fig. 2 | 4 |
| Supplementary Fig. 3 | 5 |
| Supplementary Fig. 4 | 6 |
| Supplementary Fig. 5 | 7 |
| Supplementary Fig. 6 | 8 |
| Supplementary Fig. 7 | 11 |
| Supplementary Fig. 8 | 12 |
| Supplementary Fig. 9 | 13 |
| Supplementary Fig. 10 | 14 |
| Supplementary Fig. 11 | 15 |
| Supplementary Fig. 12 | 16 |
| Supplementary Fig. 13 | 17 |
| Supplementary Fig. 14 | 18 |
| Supplementary Fig. 15 | 19 |
| Supplementary Fig. 16 | 20 |
| Supplementary Fig. 17 | 21 |
| Supplementary Fig. 18 | 23 |
| Supplementary Table. 1 | 24 |
| Supplementary Table. 2 | 26 |
| Supplementary Table. 3 | 29 |

**a**

| Antibody ID | Test | SARS | WT | Alpha | Beta | Gamma | Delta | BA.1 | BA.2 | BA.4/5 | BF.7 | BQ.1 | BQ.1.1 | XBB | XBB.1 | XBB.1.5 | CH.1.1 |
|---|---|---|---|---|---|---|---|---|---|---|---|---|---|---|---|---|---|
| 6H2 Fab | CFS | 29.73 | 3.07 | 2.68 | 4.12 | 1.88 | 5.79 | 0.09 | 0.11 | 0.01 | 0.00 | 0.01 | 0.01 | 0.00 | 0.00 | 0.00 | 0.00 |
| | neut | 0.153 | 0.881 | 1.230 | 1.115 | 1.053 | 0.887 | >100 | >100 | >100 | >100 | >100 | >100 | >100 | >100 | >100 | >100 |

**b**

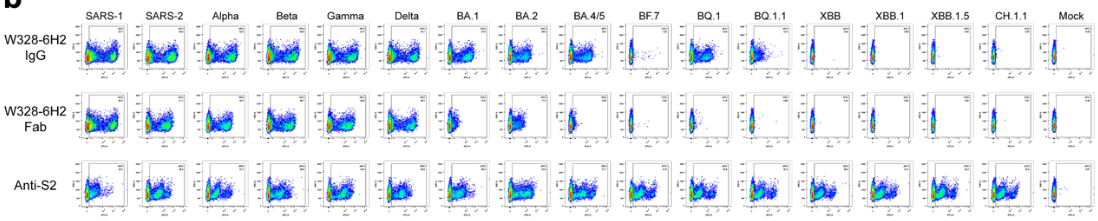


**Supplementary Fig. 1, related to Figure 1a. a** the cross-binding ability and neutralizing potency (IC50) of 6H2 Fab to SARS and SARS-CoV-2 variants carried out by cell surface staining (CFS) and neutralization (neut) experiments. For cell surface staining, the diverse S of SARS and SARS-CoV-2 variants were expressed on the surface of HEK293T cells, stained with 6H2 Fab, and detected by flow cytometry. Total fluorescence intensity (TFI) of positively stained cells were calculated and normalized to represent the relative binding ability. For neutralization, the detection limit (the highest antibody concentration used for neutralization) was 100 µg/mL. Results of both experiments were derived from two independent repeats. **b** The Gating strategies and flow cytometry results of CFS of 6H2 IgG and Fab. Anti-S2, a broadly reactive S2-specific antibody, was used as a positive control and for expression normalization. The numbers at the upper-right hand corners within each gate represent the percent of cells detected by each antibody. The Result was a representative of two independent experiments.

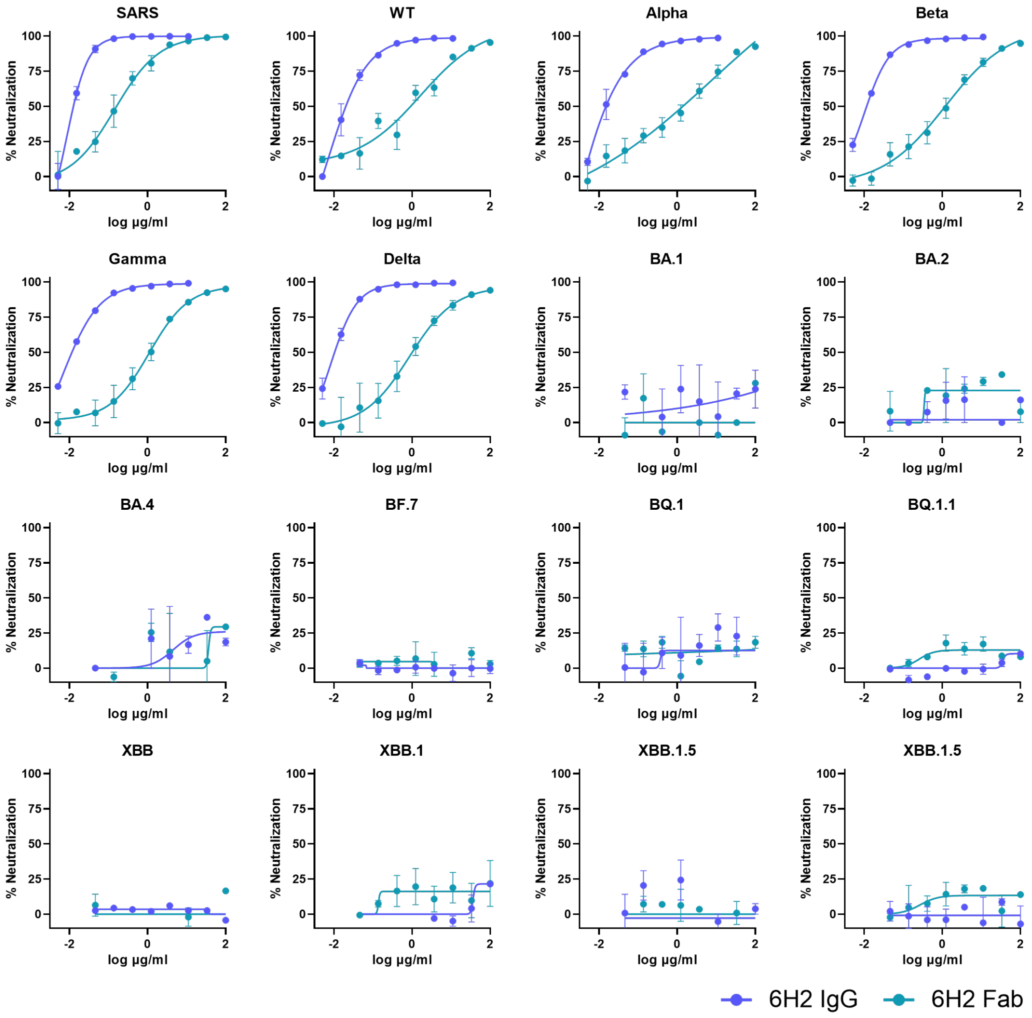


**Supplementary Fig. 2, related to Figure 1a. The neutralizing curve of IgG and Fab forms of antibody 6H2 against the panel of 16 coronaviruses with mean and s.e.m. labelled, from which the IC50 was estimated and presented in Figure 1a.** Results were derived from two independent experiments, and each included two technical replicates.

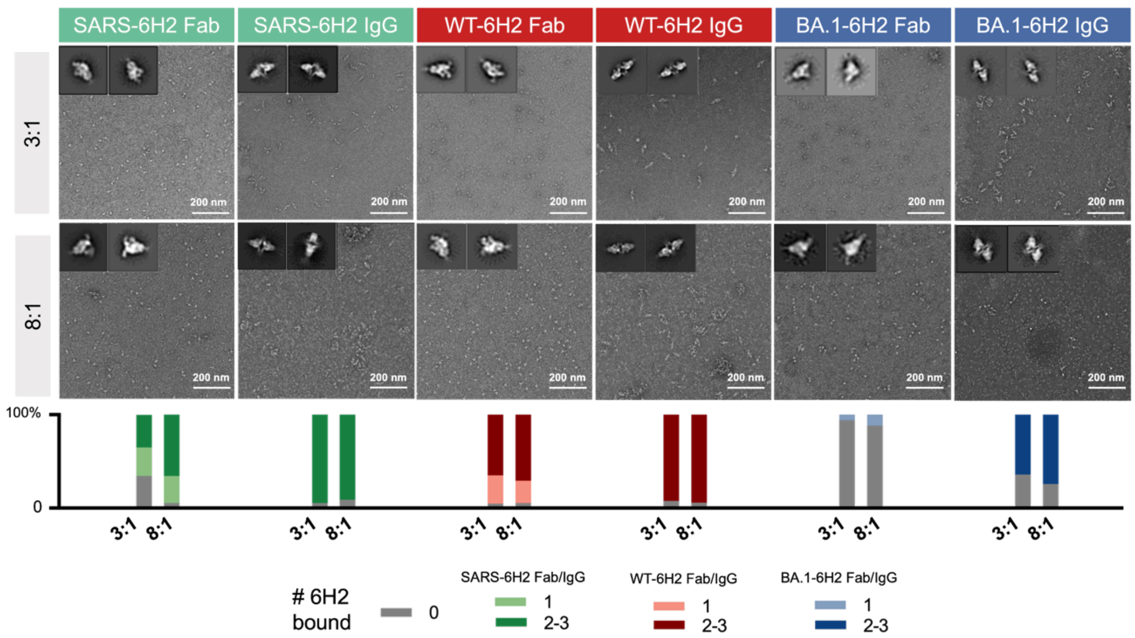


**Supplementary Fig. 3, related to Figure 1b. The effect of different ratio of Fab to protomer on binding percentage and binding mode.** SARS, WT and BA.1 S were complexed with 3-fold (~5.1 μM Fab to ~1.7 μM S protomer) and 8-fold (~13.6 μM Fab to ~1.7 μM S protomer) excess of 6H2 Fab or IgG and incubated at RT for 6h. Top: Raw micrographs and representative 2D class from nsEM analysis of 6H2 Fab or IgG complexed with S of SARS (green), WT (red) and BA.1 (blue). Bottom: NsEM semi-quantitative epitope occupancy analysis denoting the proportion of S trimers with 0, 1 and 2~3 6H2 fabs bound.

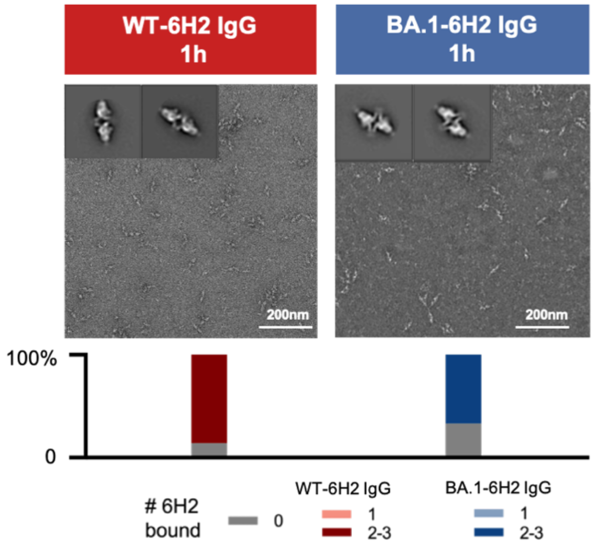


**Supplementary Fig. 4, related to Figure 1b. The effect of different incubation time on binding percentage and binding mode.** WT and BA.1 S were complexed with 3-fold (~5.1 µM Fab to ~1.7 µM S protomer) excess of 6H2 IgG and incubated at RT for 1h. Top: Raw micrographs and representative 2D class from nsEM analysis of 6H2 IgG complexed with S of WT (red) and BA.1 (blue). Bottom: Ns-EM semi-quantitative epitope occupancy analysis denoting the proportion of S trimers with 0, 1 and 2~3 6H2 fabs bound.

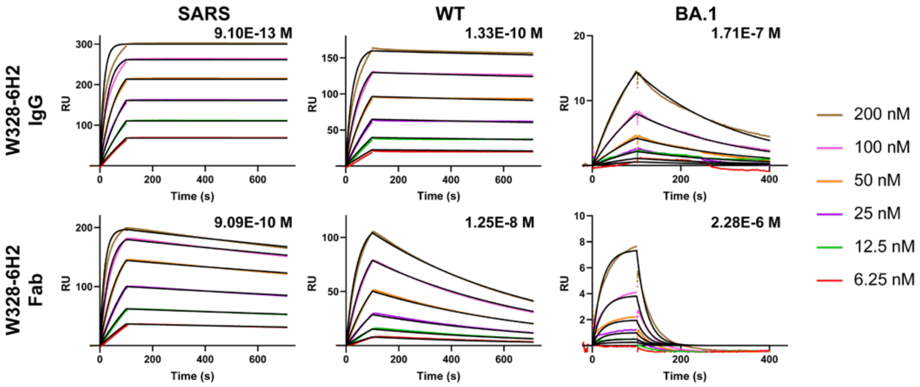


| | | SPR Binding Kinetics | | |
|---|---|---|---|---|
| | | SARS | WT | BA.1 |
| 6H2 IgG | ka (1/Ms) | 3.42E+05 | 1.95E+05 | 2.40E+04 |
| | kd (1/s) | 3.11E-07 | 2.60E-05 | 4.10E-03 |
| | KD (M) | 9.10E-13 | 1.33E-10 | 1.71E-07 |
| | R2 | 0.987 | 0.989 | 0.996 |
| 6H2 Fab | ka (1/Ms) | 2.90E+05 | 1.24E+05 | 1.94E+04 |
| | kd (1/s) | 2.63E-04 | 1.55E-03 | 4.44E-02 |
| | KD (M) | 9.09E-10 | 1.25E-08 | 2.28E-06 |
| | R2 | 0.952 | 0.953 | 0.900 |

**Supplementary Fig. 5, related to Figure 1a. Binding kinetics of 6H2 IgG and Fab to SARS, WT or BA.1 S measured by SPR.** Three types of S were immobilized on a CM5 sensor chip and serial concentrations of either IgG or Fab form of 6H2 was flowed through the system. Colored lines indicate the experimentally derived curves. Black lines represent the best fitted curves based on the experimental data. The calculated ka, kd, KD and $R^2$ values of each pair of antibody and S are shown in the bottom. The disassociation rate constant for 6H2 IgG to SARS S should be cautiously interpreted for it is below the detection limit.

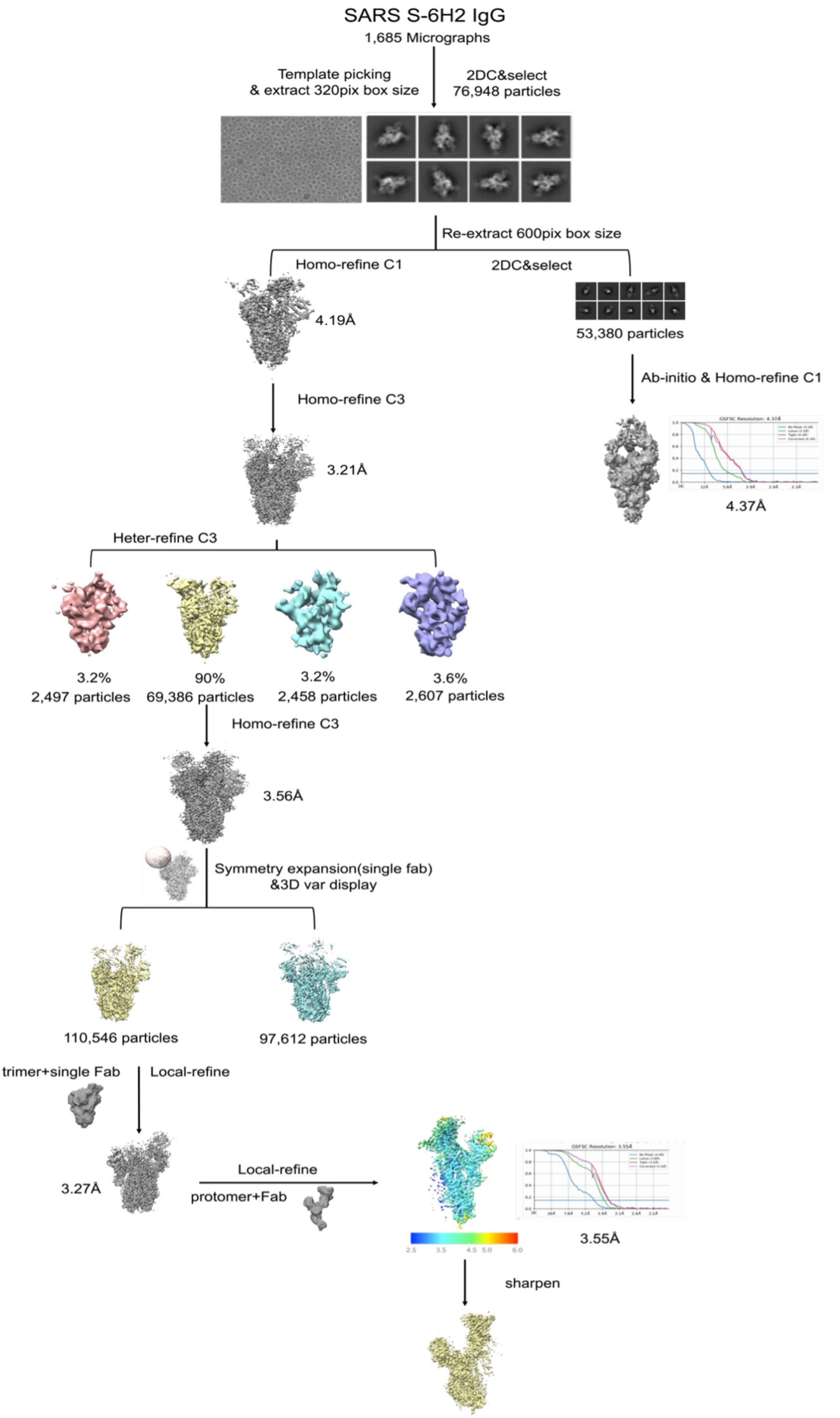
SARS S-6H2 IgG
1,685 Micrographs
Template picking
& extract 320pix box size
2DC&select
76,948 particles
Re-extract 600pix box size
Homo-refine C1
2DC&select
4.19Å
53,380 particles
Ab-initio & Homo-refine C1
Homo-refine C3
3.21Å
4.37Å
Heter-refine C3
3.2%
2,497 particles
90%
69,386 particles
3.2%
2,458 particles
3.6%
2,607 particles
Homo-refine C3
3.56Å
Symmetry expansion(single fab)
&3D var display
110,546 particles
97,612 particles
trimer+single Fab
Local-refine
3.27Å
Local-refine
protomer+Fab
3.55Å
sharpen

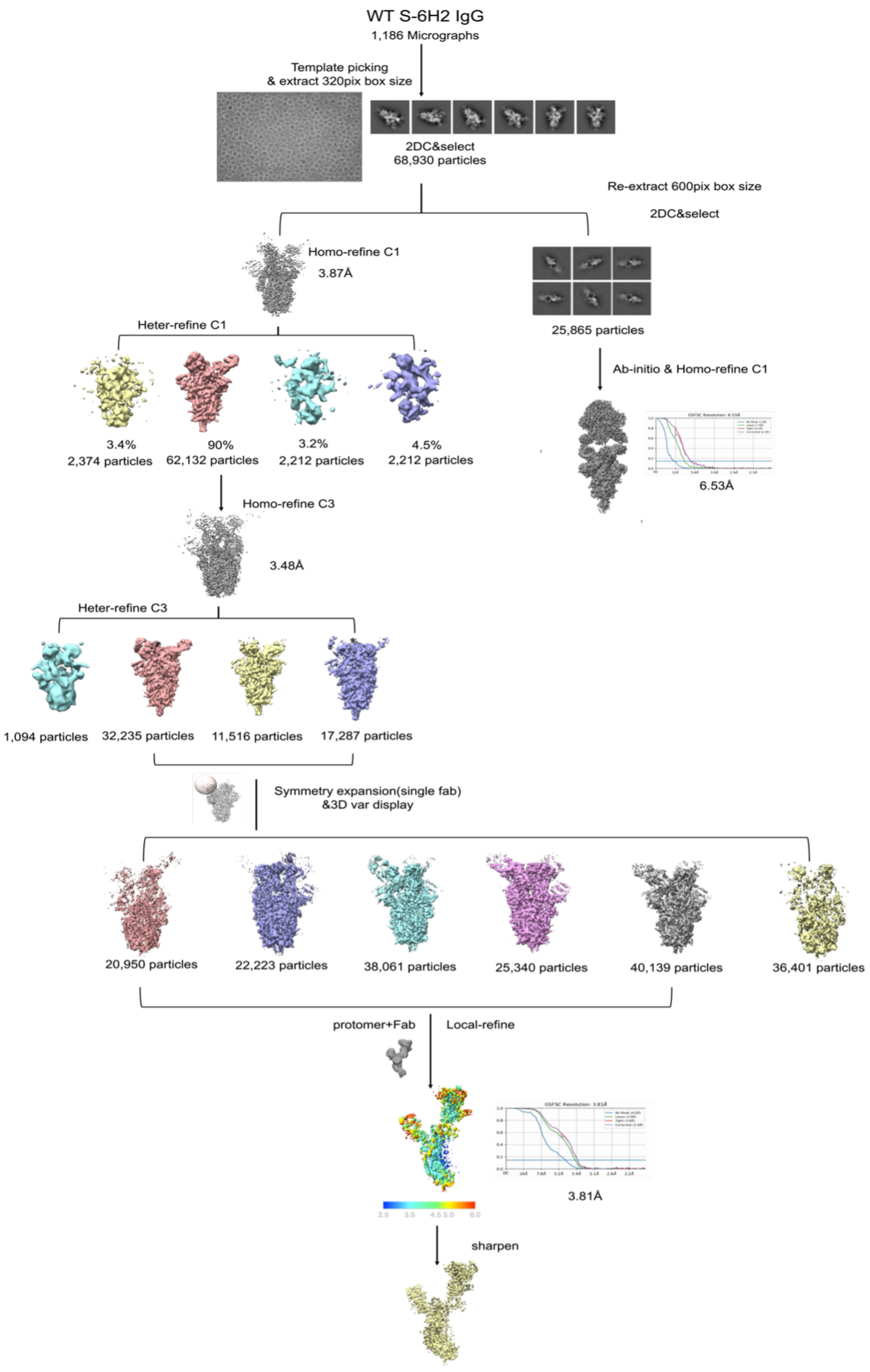
WT S-6H2 IgG
1,186 Micrographs
Template picking
& extract 320pix box size
2DC&select
68,930 particles
Re-extract 600pix box size
2DC&select
Homo-refine C1
3.87Å
25,865 particles
Heter-refine C1
3.4%
2,374 particles
90%
62,132 particles
3.2%
2,212 particles
4.5%
2,212 particles
Ab-initio & Homo-refine C1
6.53Å
Homo-refine C3
3.48Å
Heter-refine C3
1,094 particles
32,235 particles
11,516 particles
17,287 particles
Symmetry expansion(single fab)
&3D var display
20,950 particles
22,223 particles
38,061 particles
25,340 particles
40,139 particles
36,401 particles
protomer+Fab
Local-refine
3.81Å
sharpen

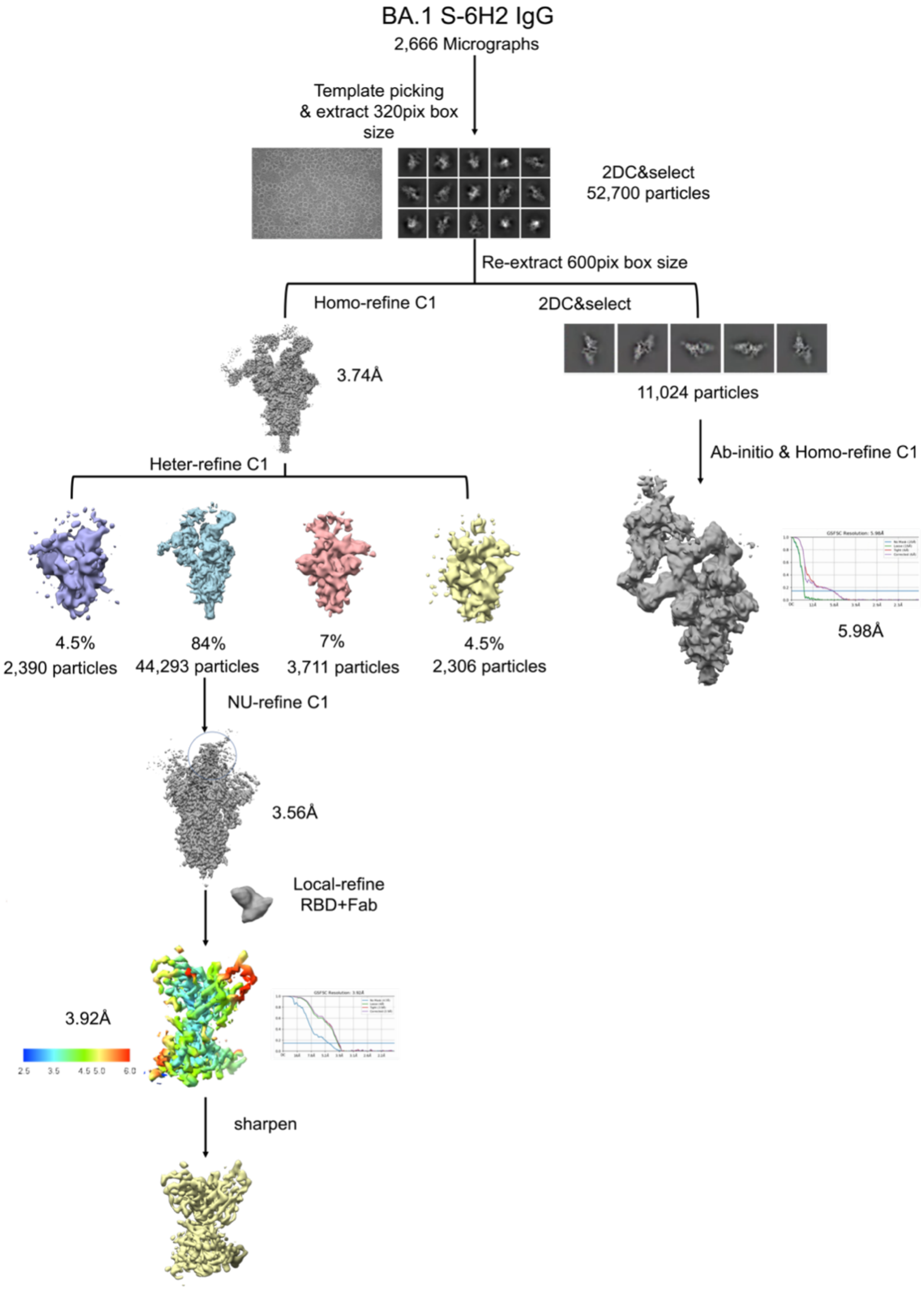


**Supplementary Fig. 6, related to Figure 1c. Data processing workflow for the SARS S-6H2 IgG, WT S-6H2 IgG and BA.1 S-6H2 IgG**. Representative raw micrographs and the reference-free 2D class averages are also presented. Local resolution evaluation of the protomer and single Fab has been shown and resolution assessment by GSFSC at 3.55 Å, 3.76 Å and 3.80 Å for SARS S-6H2 IgG, WT S-6H2 IgG and BA.1 S-6H2 IgG, respectively.

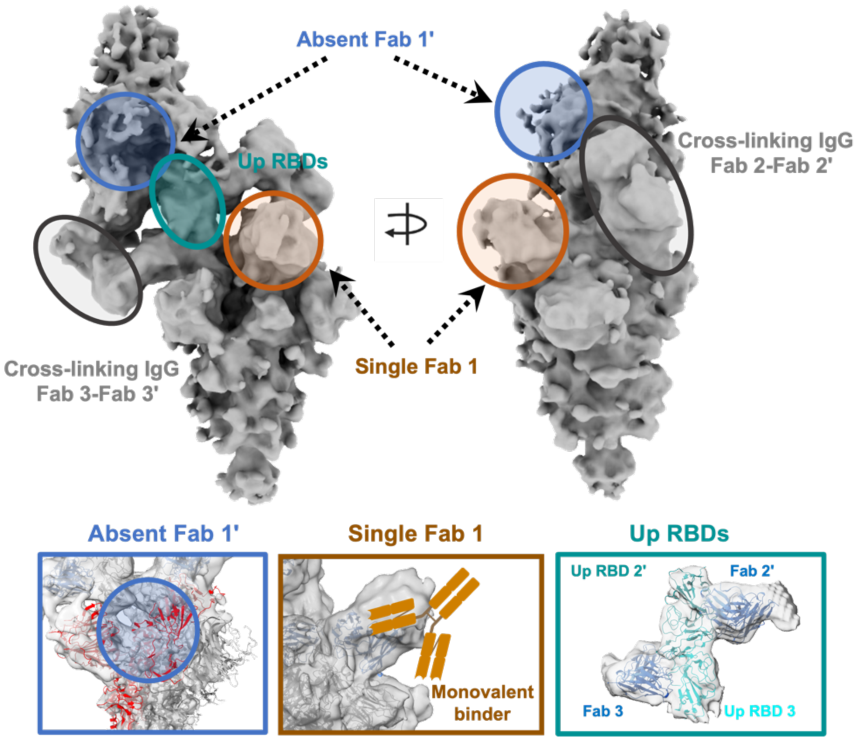


**Supplementary Fig. 7, related to Figure 1c. The global conformation of offset head-to-head dimer-trimer of BA.1-S 6H2 IgG complex achieved by cryo-EM and processed in a 600 Å box size.** A total of two IgGs each cross-linked a “down” RBD and an “up” RBD from the opposite S. The third RBD of both S did not form the cross-linking binding mode with one binding to a single Fab arm of 6H2 IgG (Single Fab1) and the other being unoccupied (Absent Fab1’).

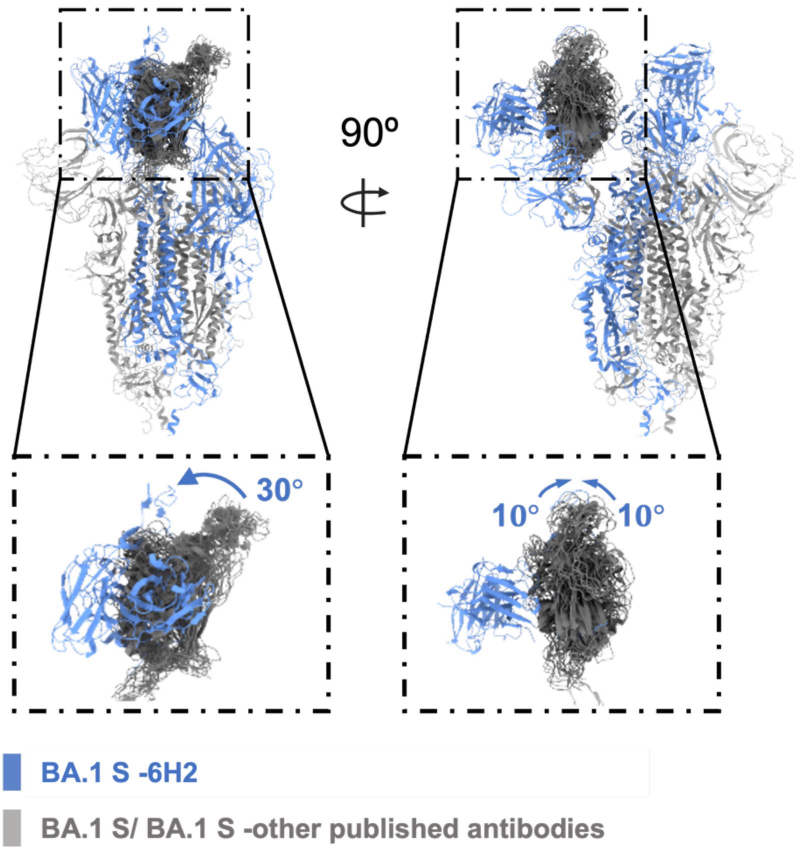


**Supplementary Fig. 8, related to Figure 2d. Docking the published PDB of BA.1 S and antibody-BA.1 complexion.** The angle of rotated from two directions has been labeled. Docked pdb: BA.1 S-open(7wvn), JMB2002-BA.1(7xod), MB.02-BA.1(8dzi), 35B5-BA.1(7wly), S3H3-BA.1(7wk9), XGv282-BA.1(7we7), XGv347-BA.1(7wea), S309-BA.1(7xco), Bn03-BA.1(7whk), A19-46.1-BA.1(7tca).

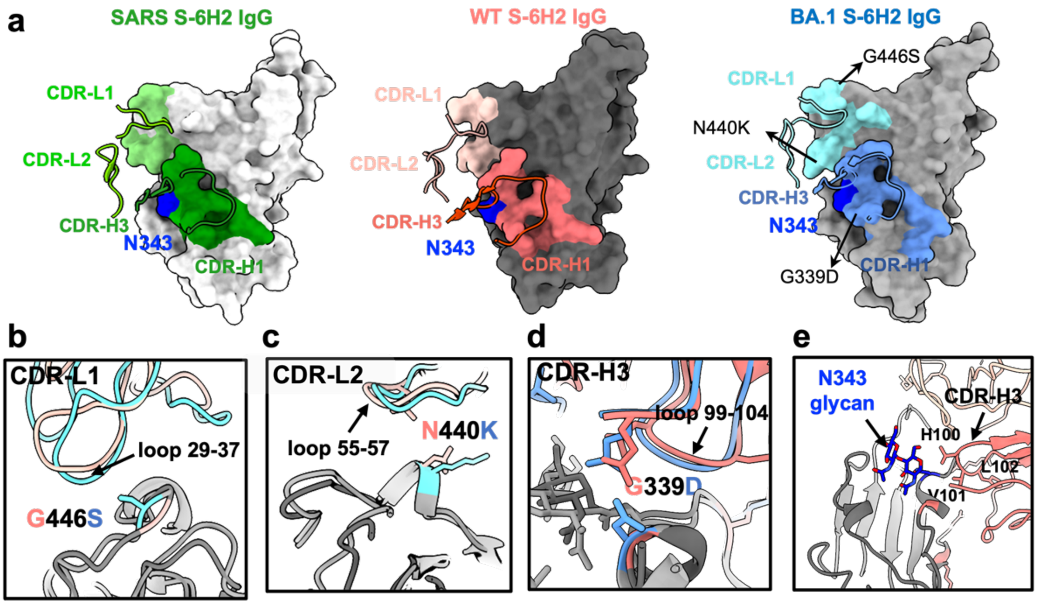


**Supplementary Fig. 9, related to Figure 3a. The structure analysis of SARS S-6H2 IgG, WT S-6H2 IgG and BA.1 S-6H2 IgG.** **a** The footprint of 6H2 IgG on the RBD region of SARS S (Flora and Forest green, for L and H chain), WT S(Light pink and Salmon) and BA.1 S (Sky blue and Cornflower blue). Mutations from BA.1 S have been labeled. **b-d** The detailed interactions of CDR-L1 (b), CDR-L2 (c) and CDR-H3 (d) of 6H2 with RBD region of SARS, WT and BA.1 S. **e** Residues involved in interactions between 6H2 and WT S glycoprotein (N343).

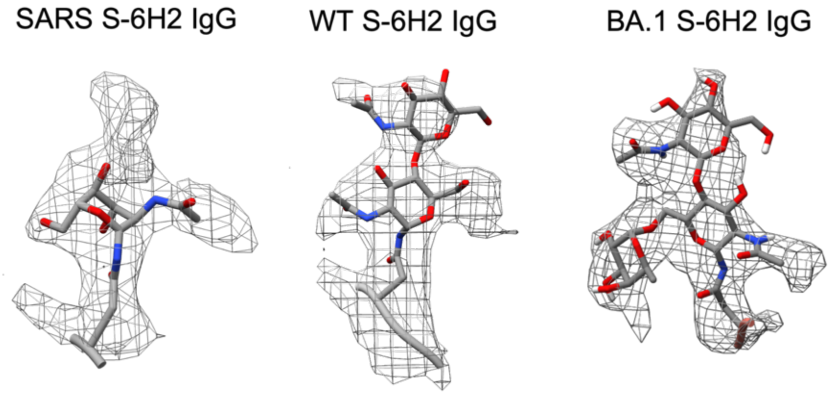


**Supplementary Fig. 10, related to Figure 3a. Density resolving the N330/N343 glycan.** The sharpened cryo-EM maps are rendered as mesh with the corresponding model shown as sticks colored by element. The N330 and N343 corresponds to the cryo-EM map.

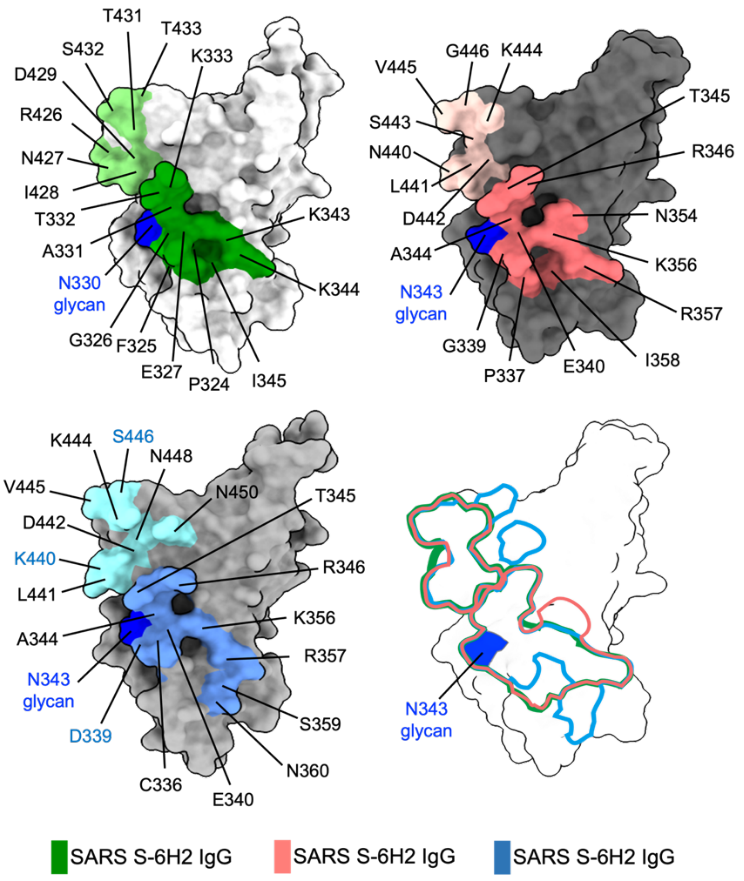


**Supplementary Fig. 11, related to Figure 3a. The epitope of SARS S-6H2 IgG (green), WT S-6H2 IgG (red) and BA.1 S-6H2 IgG (blue)**. All the contact residues have been labeled and the mutations of BA.1 have been colored by blue. Overlapping of the footprint shows the almost identical epitopes.

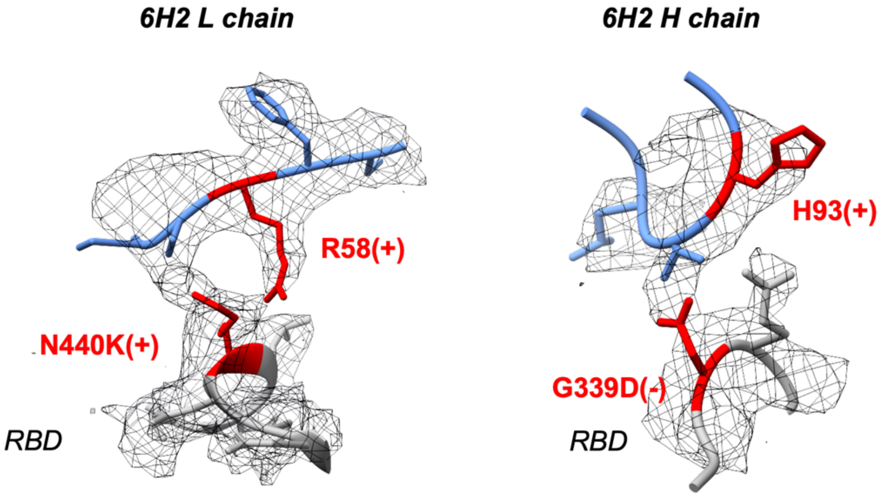


**Supplementary Fig. 12, related to Figure 3a. Density resolving the G339D and N440K mutations in BA.1 S.** The sharpened cryo-EM map (6H2 Fab and RBD region) as rendered as mesh with the corresponding model shown as sticks (6H2 Fab: blue; RBD: grey; Mutations: red).

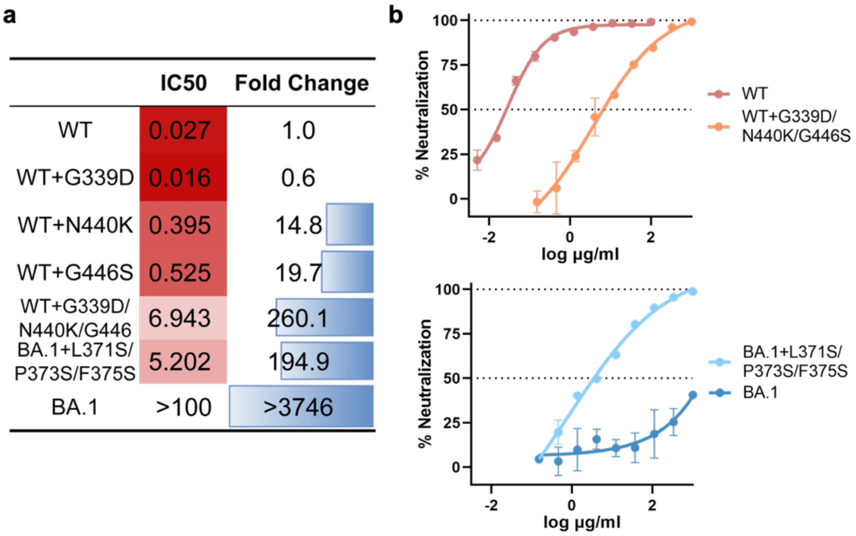


**Supplementary Fig. 13, related to Figure 3b-e.** a. IC50 calculated from the fitted neutralization curves in Fig 3c&e with the fold changes presented as a bar chart. b. Repeated neutralization of 6H2 against WT, WT with G339D-N440K-G446S triple-site mutations (WT-M), BA.1 and BA.1 with L371S-P373S-F375S triple-site mutations (BA.1-M) to validate the neutralization efficacy of 6H2 IgG. The antibody concentration was added up to 1000 µg/mL before serial dilution against WT-M, BA.1-M and BA.1. For reference, neutralization against WT was also tested with the highest concentration of 10 µg/mL. The neutralizing curves against the 4 variants were drawn from the repeated data.

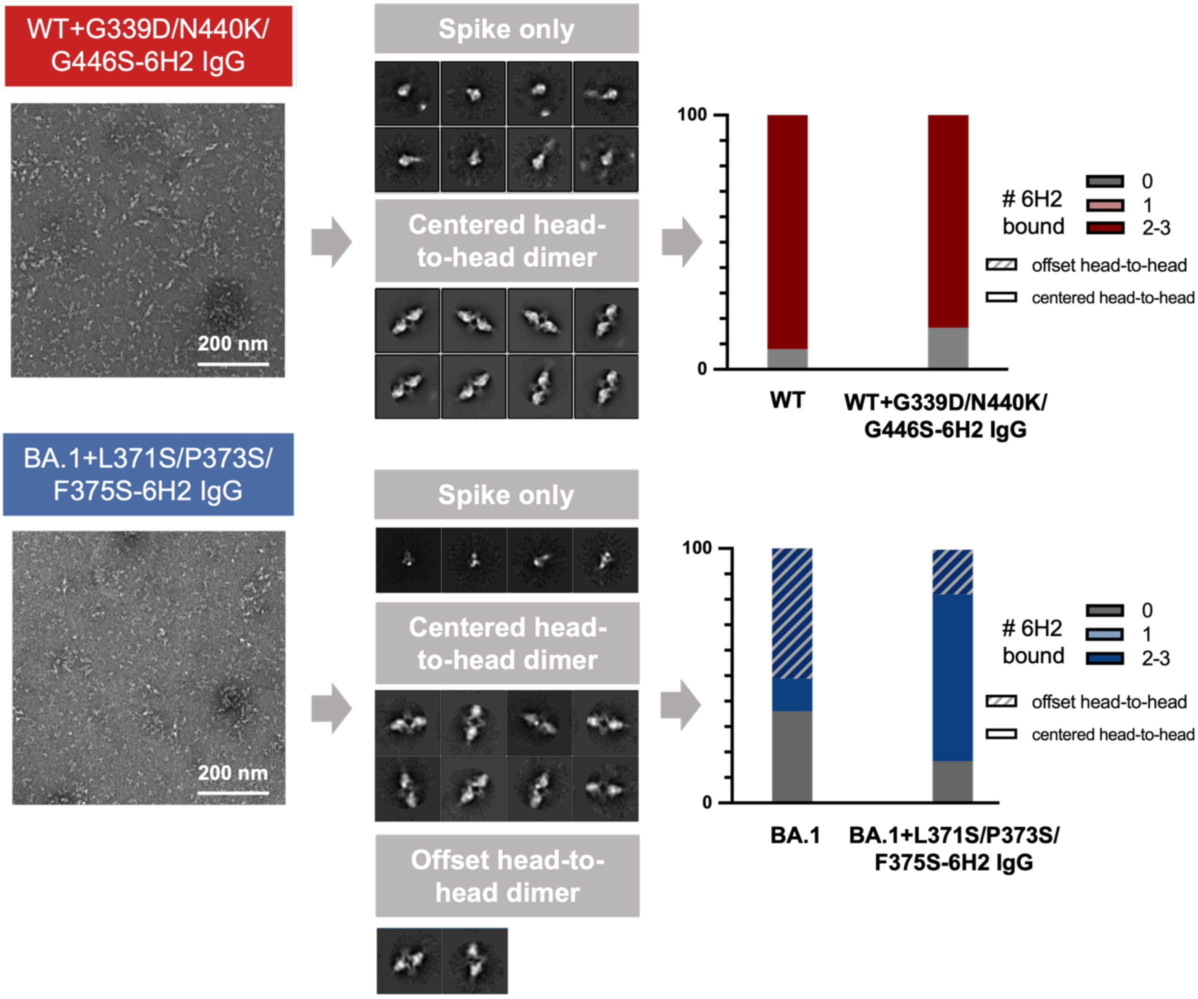


**Supplementary Fig.14, related to Figure 3b-e. Ns-EM analysis on WT- M S-6H2 IgG and BA.1-M S-6H2 IgG.** WT-M and BA.1-M S were complexed with 3-fold (~5.1 µM Fab to ~1.7 µM S protomer) excess of 6H2 IgG and and incubated at RT for 6h. Left: Raw micrographs and representative 2D class from nsEM analysis of 6H2 IgG complexed with S of WT-M (red) and BA.1-M (blue). Right: NsEM analysis denoting the proportion of S trimers with 0, 1 and 2~3 6H2 fabs bound and distinct dimer-trimer structures.

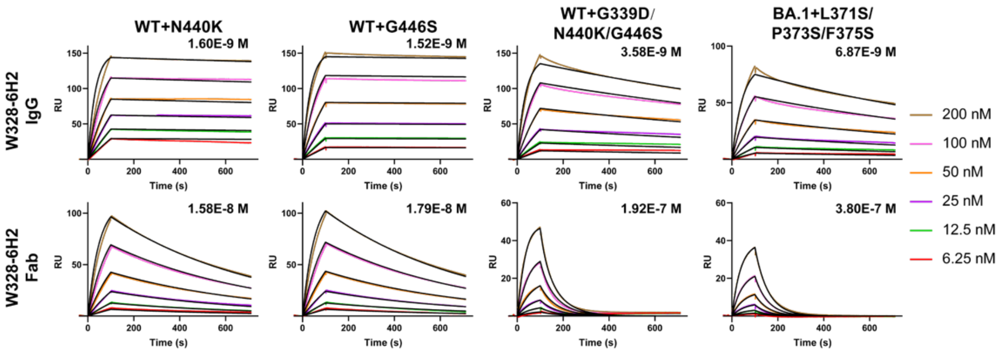


| | | SPR Binding Kinetics | | | |
|---|---|---|---|---|---|
| | | WT+N440K | WT+G446S | WT+G339D/ N440K/G446 | BA.1+L371S/ P373S/F375S |
| 6H2 IgG | ka (1/Ms) | 1.44E+05 | 1.50E+05 | 1.40E+05 | 1.06E+05 |
| | kd (1/s) | 2.30E-04 | 2.28E-04 | 5.02E-04 | 7.25E-04 |
| | KD (M) | 1.60E-09 | 1.52E-09 | 3.58E-09 | 6.87E-09 |
| | R2 | 0.940 | 0.951 | 0.937 | 0.943 |
| 6H2 Fab | ka (1/Ms) | 9.72E+04 | 8.94E+04 | 3.45E+05 | 4.63E+04 |
| | kd (1/s) | 1.54E-03 | 1.60E-03 | 6.62E-02 | 1.76E-02 |
| | KD (M) | 1.58E-08 | 1.79E-08 | 1.92E-07 | 3.80E-07 |
| | R2 | 0.940 | 0.942 | 0.942 | 0.968 |

**Supplementary Fig. 15, related to Figure 3b-e. Binding kinetics of 6H2 IgG and Fab to WT with N440K and G446S single-site mutation and WT-M mutation and BA.1-M mutation by SPR.** Four types of S were immobilized on a CM5 sensor chip and serial concentrations of either IgG or Fab form of 6H2 was flowed through the system. Colored lines indicate the experimentally derived curves. Black lines represent the best fitted curves based on the experimental data. The calculated ka, kd, KD and $R^2$ values of each pair of antibody and S are shown in the bottom.

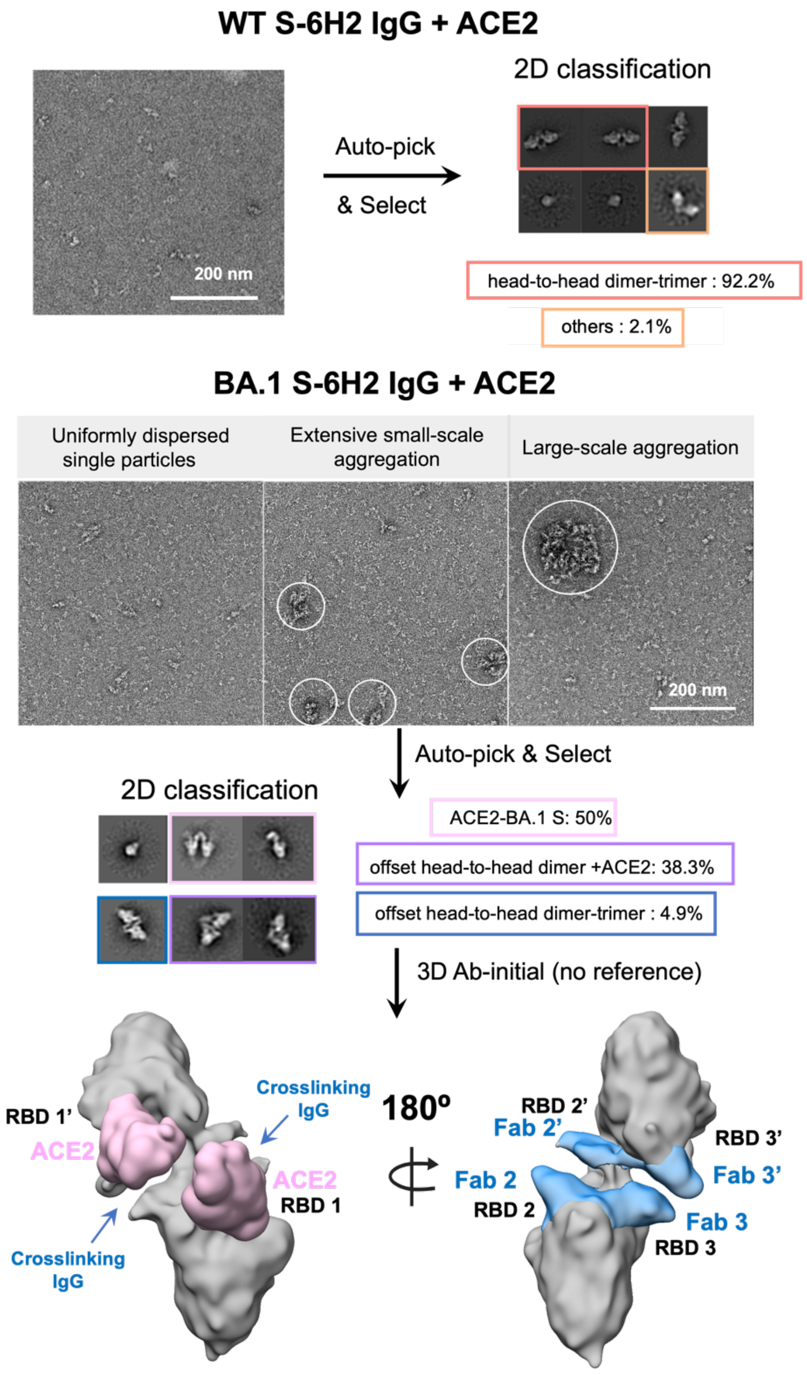


**Supplementary Fig. 16, related to Figure 3a-c. Raw micrographs, 2D class average and 3D Ab-initial map of WT or BA.1 S-6H2 complex added ACE2 applied by ns-EM.** WT or BA.1 S were complexed with 6H2 IgG at 3× molar excess of Fab/protomer ratio and incubated incubated at RT for 5 hours. Then ACE2 was added at 3× molar excess of ACE2/protomer ratio and incubated incubated at RT overnight. Raw micrographs, 2D class average as well as proportion of each class calculated from ns-EM 2DC have been shown. For BA.1 S-6H2-ACE2 complex, as side views of 3D reconstructions are shown in the bottom and antibodies or ACE2 are colored according to those initially designated in Fig. 4c.

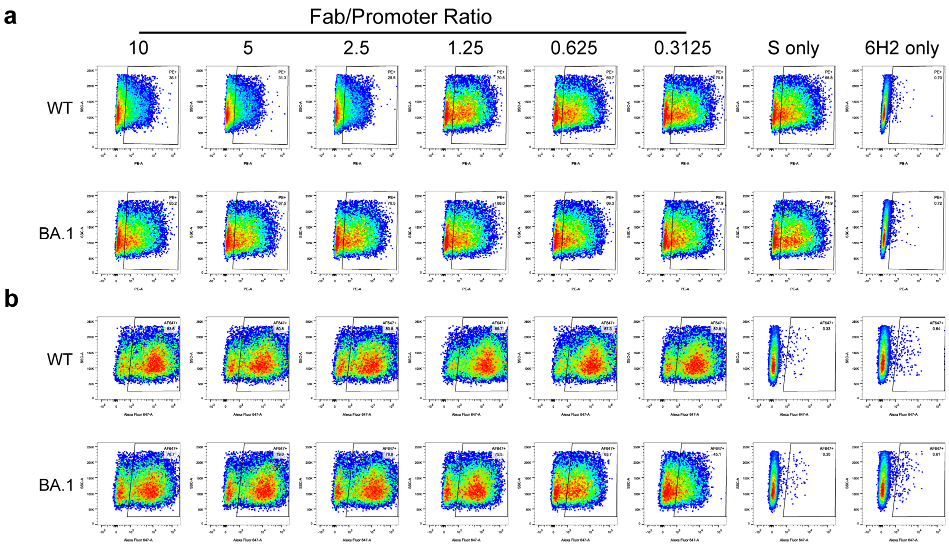


**Supplementary Fig. 17, related to figure 4g-h. The Gating strategies and flow cytometry results of CFS of S-6H2 IgG complexes.** 293T-ACE2 cells were incubated with WT/BA.1 S-6H2 IgG of different Fab/Protomer ratio, while cells incubated with Spike or 6H2 alone were used as controls. Spike(**a**) or 6H2(**b**) attached to the cell surface were fluorescence labelled. The numbers at the upper-right hand corners within each gate indicate the percentage of positive cells detected. The result was a representative of two independent experiments.

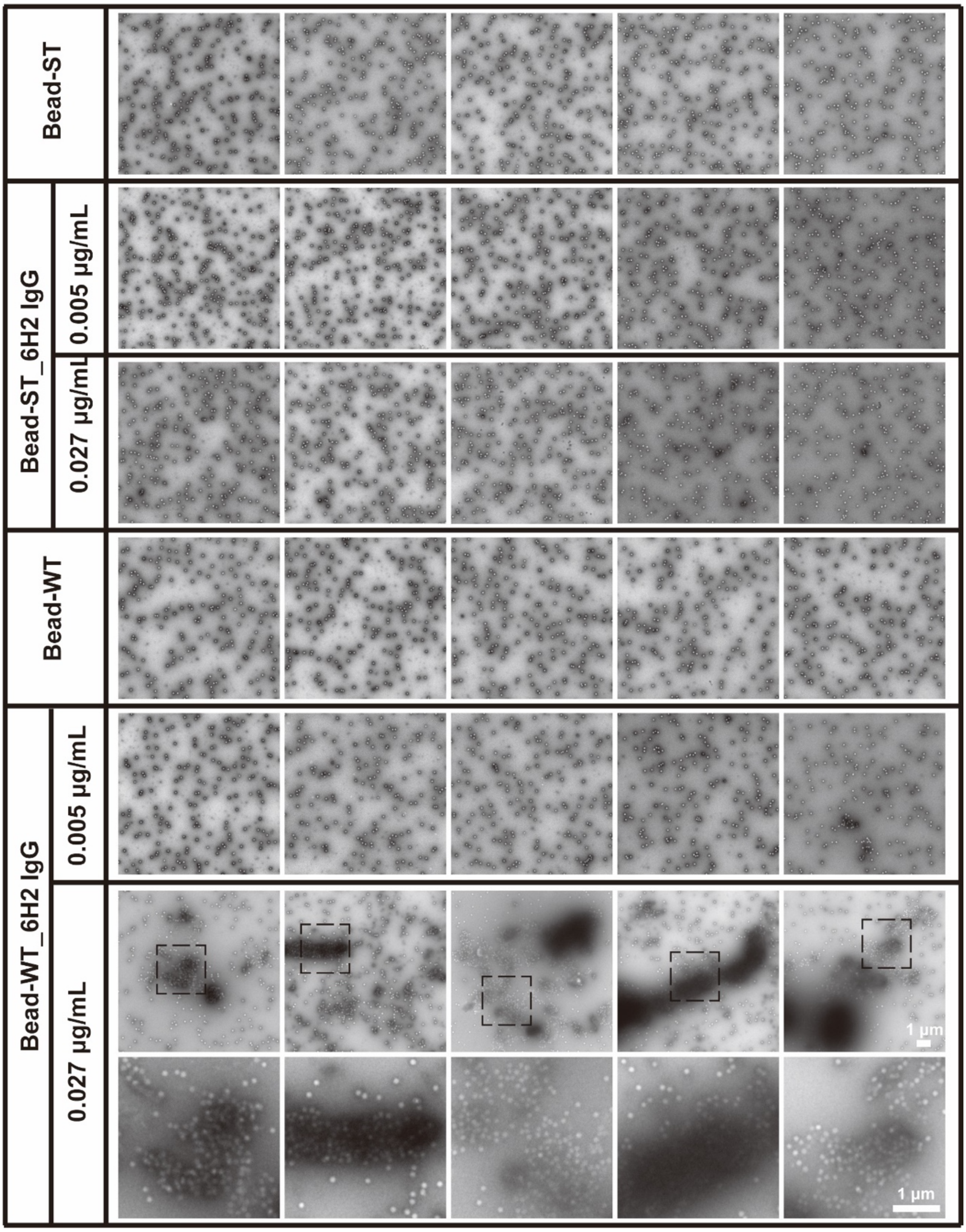
Bead-ST
Bead-ST_6H2 IgG
0.005 μg/mL
0.027 μg/mL
Bead-WT
Bead-WT_6H2 IgG
0.005 μg/mL
0.027 μg/mL
1 μm
1 μm

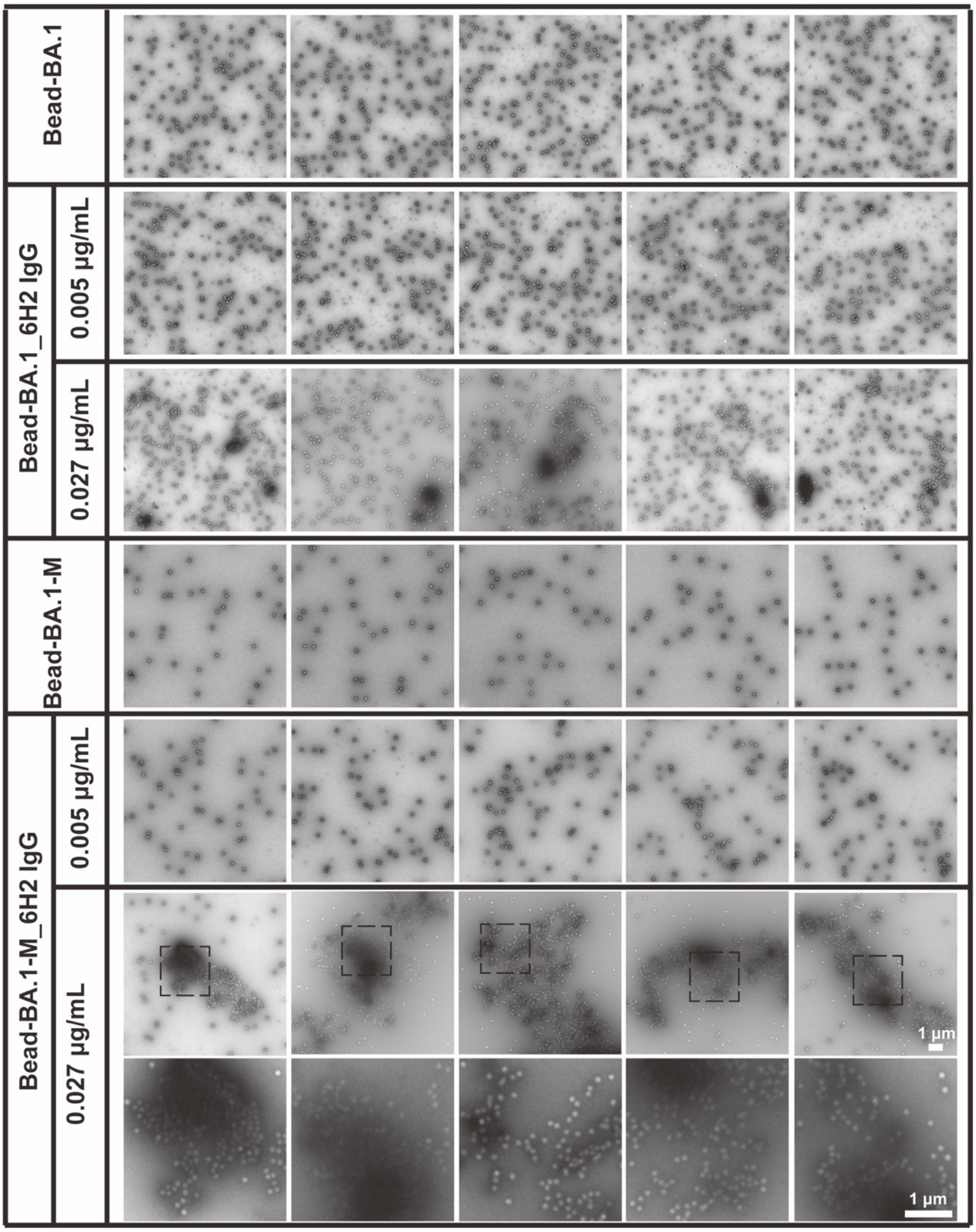


**Supplementary Fig. 18, related to Figure 5b. Representative raw micrographs characterizing the distribution patterns of beads modified with WT, BA.1 or BA.1-M S following the addition of 6H2 IgG at different concentrations.** The Streptactin-coated beads and either the WT, BA.1 or BA.1-M S are blended and incubated at RT for 4 hours without any further washing steps. 6H2 IgG at varying concentrations (0.005 μg/mL, 0.027 μg/mL) was introduced into the WT/BA.1/BA.1-M-beads solution and incubated at RT for 1 hour. Finally, the beads suspensions were diluted with TBS and deposited on a carbon-coated electron microscope grid, followed by negatively staining with 2% uranyl acetate. The distribution states of beads were performed on TEM. Scale bar: 1 μm. The scale bar in the last image is applicable to all other images in the same panel.

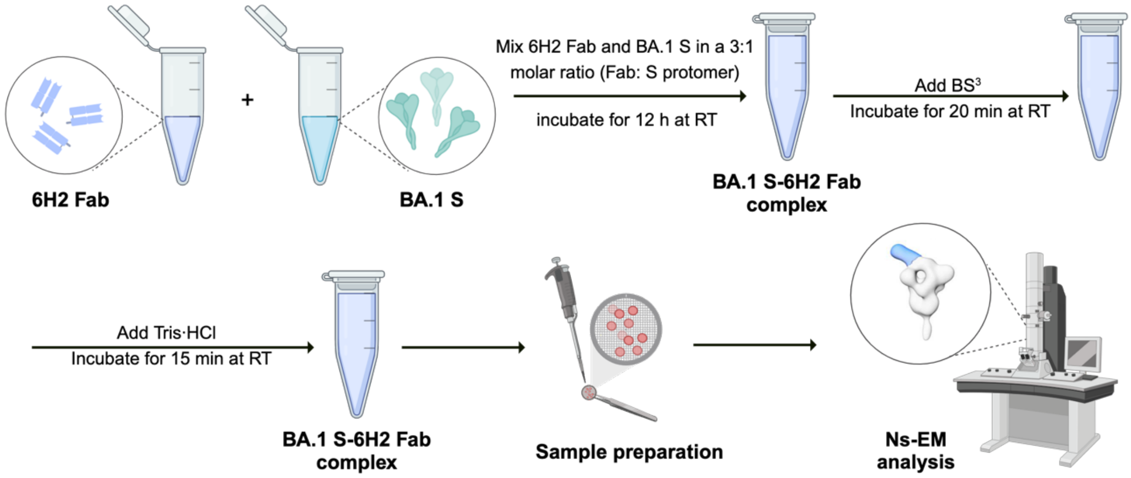


**Supplementary Fig. 19, related to Figure 1b. Procedure of $BS^3$ Crosslinker.** BA.1 S and 6H2 Fab were first mixed in a 3:1 molar ratio (Fab: S protomer) in the buffer (pH 7.5) containing 20 mM HEPES and 150 mM NaCl. After a 12-hour incubation at RT, the BA.1 S-6H2 Fab complex was crosslinked with 0.5 mM $BS^3$ for 20 min. The crosslinker reaction was subsequently quenched by adding 400 mM Tris (pH 8) to achieve a final concentration of 100 mM Tris, followed by a 15-minute incubation at RT.

**Supplementary Table 1.** SEC traces, SDS-PAGE and negative-stain EM images of purified antibodies, S proteins and ACE2. N.A.: not available.

| | Sample | FPLC | SDS-Page | Ns-EM |
|---|---|---|---|---|
| 1 | 6H2 IgG/Fab | N.A. | 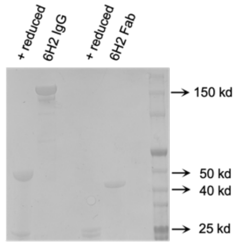 | 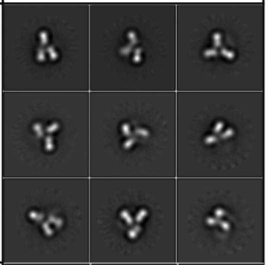 |
| 2 | SARS 2p | 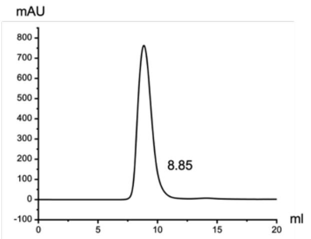 | 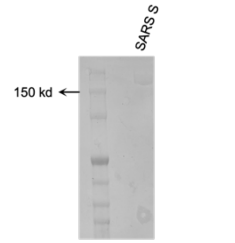 | 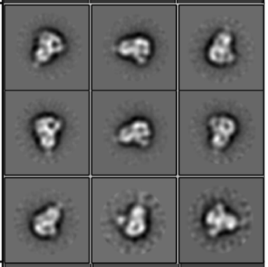 |
| 3 | WT 6p | N.A. | 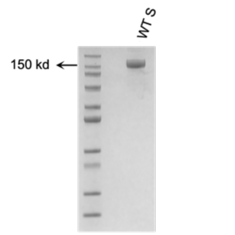 | 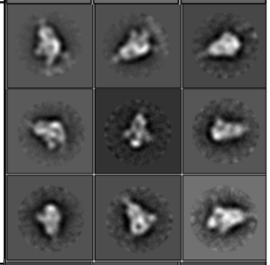 |
| 4 | Omicron BA 1 6p | 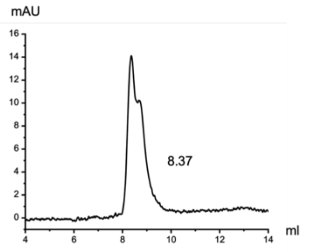 | 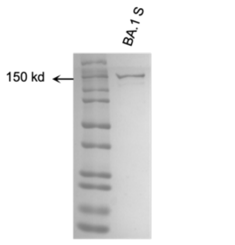 | 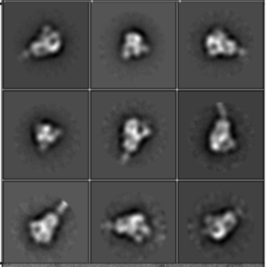 |
| 5 | Delta 2p | N.A. | 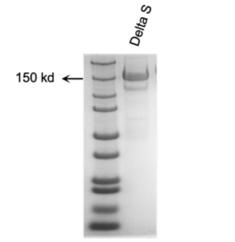 | 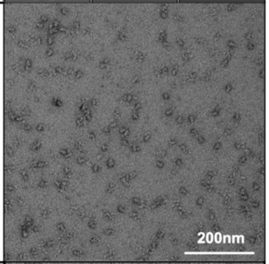 |
| 6 | WT+ mutations (N440K) | 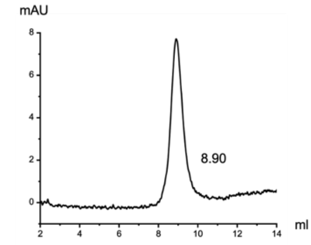 | 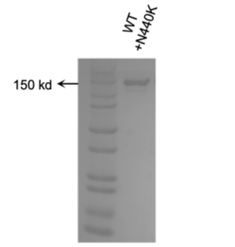 | 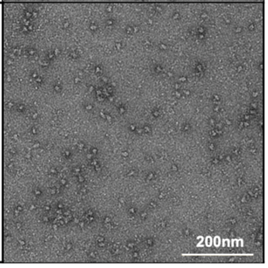 |

| | Sample | FPLC | SDS-Page | Ns-EM |
|---|---|---|---|---|
| 7 | WT+ mutations (G446S) | 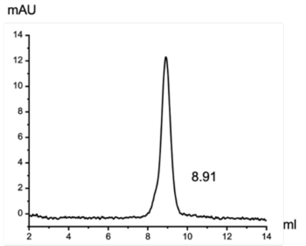  | 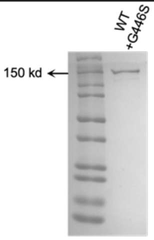  | 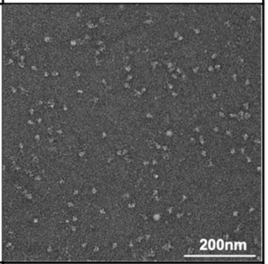  |
| 8 | WT+ Mutation (G339D/N440K/ G446S) | N.A. | N.A. | 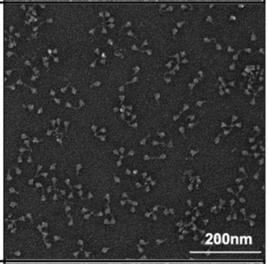  |
| 9 | BA.1+ Mutation (L371S/P373S/ F375S) | N.A. | N.A. | 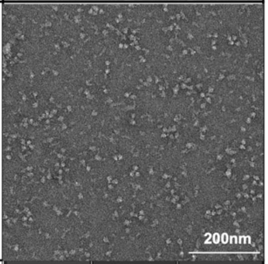  |
| 10 | ACE2 (dimer) | N.A. | N.A. | 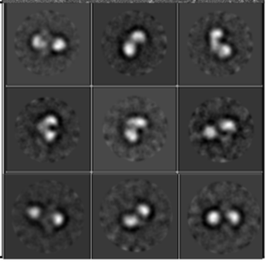 |

**Supplementary Table 2. Analysis of antibodies and spike protein complexes based on negative-stain EM.** Data tables, indicating construct names (including EMD accession numbers), mutations, observed classes and number of particles per class. N.A.: not available.

| Construct: | SARS S-6H2 IgG (Fab/protomer=3:1) EMD-36257 | |
|---|---|---|
| Class: | complex | Spike only |
| # Particles: | 5,059 | 585 |
| % Total: | 89.6% | 10.4% |

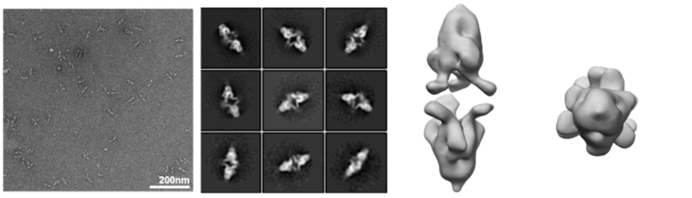


| Construct: | SARS S-6H2 Fab (Fab/protomer=3:1) EMD-35953 | |
|---|---|---|
| Class: | complex | Spike only |
| # Particles: | 4,146 | 2,232 |
| % Total: | 65.0% | 35.0% |

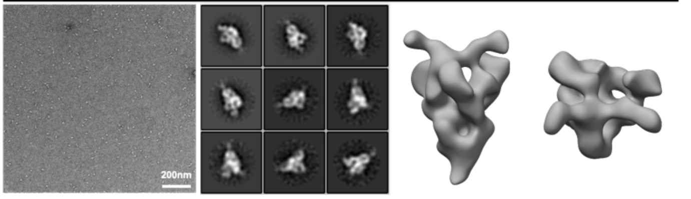


| Construct: | WT S-6H2 IgG (Fab/protomer=3:1) EMD-36267 | |
|---|---|---|
| Class: | complex | Spike only |
| # Particles: | 4,001 | 670 |
| % Total: | 85.7% | 14.3% |

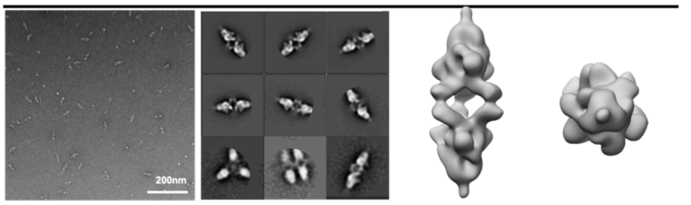


| Construct: | WT S-6H2 Fab (Fab/protomer=3:1) EMD-35961 | |
|---|---|---|
| Class: | complex | Spike only |
| # Particles: | 7,277 | 383 |
| % Total: | 95.0% | 5.0% |

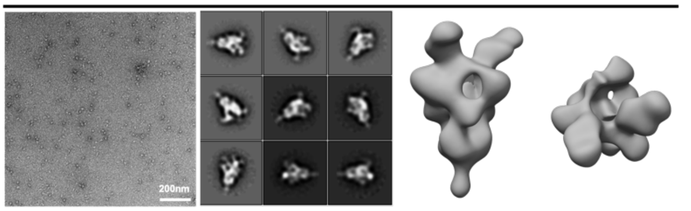


| Construct: | Delta S-6H2 IgG (Fab/protomer=3:1) | |
|---|---|---|
| Class: | complex | Spike only |
| # Particles: | 5,059 | 586 |
| % Total: | 89.6% | 10.4% |

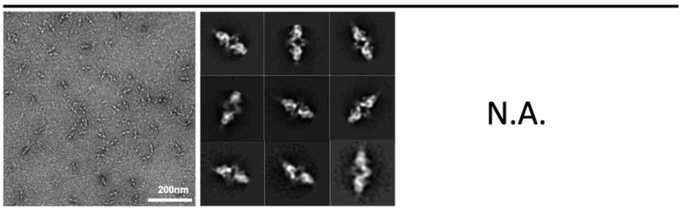


| Construct: | BA.1 S-6H2 IgG (Fab/protomer=3:1) EMD-35963 | |
|---|---|---|
| Class: | complex | Spike only |
| # Particles: | 13,429 | 15,134 |
| % Total: | 47.1% | 52.9% |

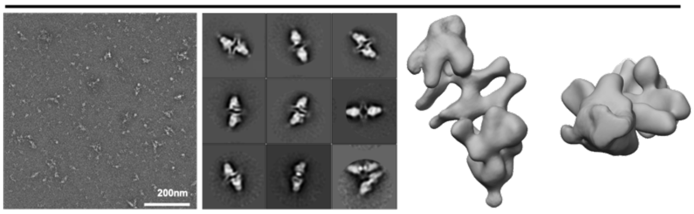


| Construct: | BA.1 S-6H2 Fab (Fab/protomer=3:1) EMD-35962 | |
|---|---|---|
| Class: | complex | Spike only |
| # Particles: | 613 | 9,693 |
| % Total: | 6.3% | 93.7% |

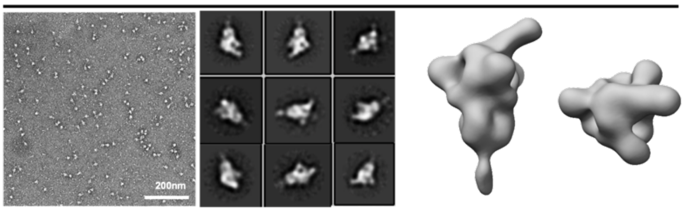

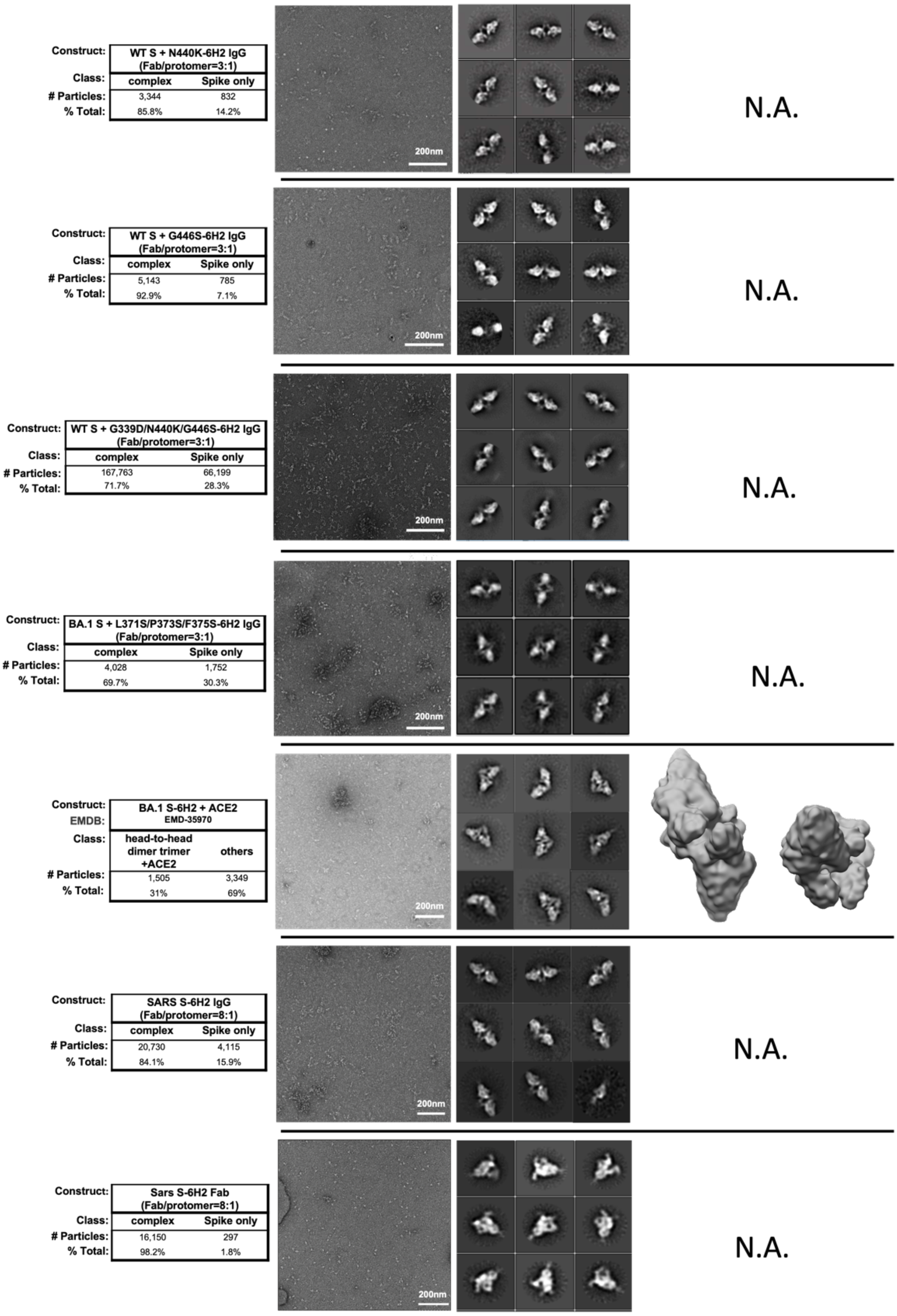
Construct: WT S + N440K-6H2 IgG (Fab/protomer=3:1)
Class: complex Spike only
### Particles: 3,344 832
% Total: 85.8% 14.2%
200nm
N.A.
Construct: WT S + G446S-6H2 IgG (Fab/protomer=3:1)
Class: complex Spike only
### Particles: 5,143 785
% Total: 92.9% 7.1%
200nm
N.A.
Construct: WT S + G339D/N440K/G446S-6H2 IgG (Fab/protomer=3:1)
Class: complex Spike only
### Particles: 167,763 66,199
% Total: 71.7% 28.3%
200nm
N.A.
Construct: BA.1 S + L371S/P373S/F375S-6H2 IgG (Fab/protomer=3:1)
Class: complex Spike only
### Particles: 4,028 1,752
% Total: 69.7% 30.3%
200nm
N.A.
Construct: BA.1 S-6H2 + ACE2
EMDB: EMD-35970
Class: head-to-head dimer trimer +ACE2 others
### Particles: 1,505 3,349
% Total: 31% 69%
200nm
Construct: SARS S-6H2 IgG (Fab/protomer=8:1)
Class: complex Spike only
### Particles: 20,730 4,115
% Total: 84.1% 15.9%
200nm
N.A.
Construct: Sars S-6H2 Fab (Fab/protomer=8:1)
Class: complex Spike only
### Particles: 16,150 297
% Total: 98.2% 1.8%
200nm
N.A.

| Construct: | WT S-6H2 IgG (Fab/protomer=8:1) | |
|---|---|---|
| Class: | complex | Spike only |
| # Particles: | 15, 573 | 1, 907 |
| % Total: | 89% | 11% |

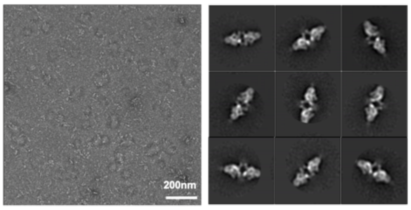


N.A.

| Construct: | WT S-6H2 Fab (Fab/protomer=8:1) | |
|---|---|---|
| Class: | complex | Spike only |
| # Particles: | 28,749 | 1,683 |
| % Total: | 94.5% | 5.5% |

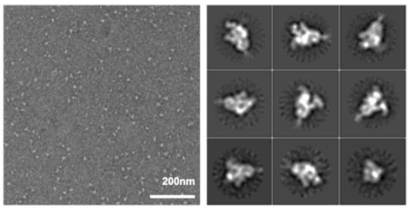


N.A.

| Construct: | BA.1 S-6H2 IgG (Fab/protomer=8:1) | |
|---|---|---|
| Class: | complex | Spike only |
| # Particles: | 6,965 | 4,174 |
| % Total: | 62.5% | 37.5% |

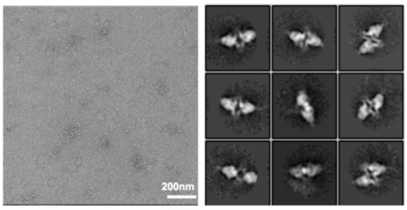


N.A.

| Construct: | BA.1 S-6H2 Fab (Fab/protomer=8:1) | |
|---|---|---|
| Class: | Complex | Spike only |
| # Particles: | 1,187 | 9,059 |
| % Total: | 11.5% | 88.5% |

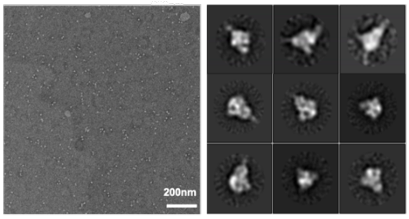


N.A.

| Construct: | WT S-6H2 IgG (Fab/protomer=3:1 1h) | |
|---|---|---|
| Class: | complex | Spike only |
| # Particles: | 32,822 | 10,751 |
| % Total: | 75.4% | 24.6% |

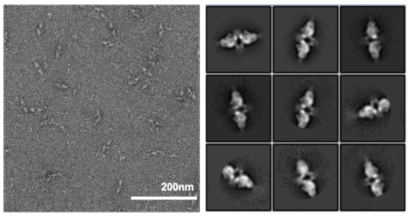


N.A.

| Construct: | BA.1 S-6H2 IgG (Fab/protomer=3:1 1h) | |
|---|---|---|
| Class: | complex | Spike only |
| # Particles: | 9,078 | 9,032 |
| % Total: | 50.2% | 49.8% |

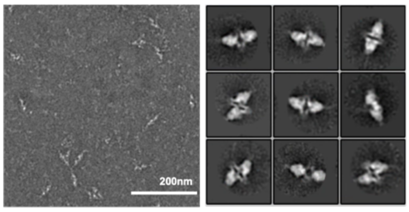


N.A.

**Supplementary Table 3.** Cryo-EM data collection, refinement and validation statistics.

**Supplementary Table 3-1.** Cryo-EM density map of SARS S-6H2 IgG, WT S-6H2 IgG and BA.1 S-6H2 IgG

| Sample | SARS S-6H2 IgG<br>EMD-35986 | WT S-6H2 IgG<br>EMD-35995 | BA.1 S-6H2 IgG<br>EMD-36058 |
|---|---|---|---|
| **Data collection and processing** | | | |
| Magnification | 29000× | 29000× | 29000× |
| Voltage (kV) | 300 | 300 | 300 |
| Electron exposure ($e^{-}/Å^2$) | 50 | 50 | 50 |
| Defocus range (μm) | -1.2 to -1.5 | -1.2 to -1.5 | -1.2 to -1.5 |
| Pixel size (Å) | 0.97 | 0.97 | 0.97 |
| Symmetry imposed | C1 | C1 | C1 |
| Final particle (no.) | 53,380 | 23,820 | 11,024 |
| Map resolution (Å) | 4.37 | 6.53 | 5.98 |

**Supplementary Table 3-2.** Local refined density maps of 6H2 Fab and RBD region of SARS S-6H2 IgG, WT S-6H2 IgG and BA.1 S-6H2 IgG

| Sample | SARS S-6H2 IgG<br>(Local refine)<br>EMD-36113<br>PDB:8JAG | WT S-6H2 IgG<br>(Local refine)<br>EMD-36121<br>PDB:8JAP | BA.1 S-6H2 IgG<br>(Local refine)<br>EMDB-36122<br>PDB:8JAM |
|---|---|---|---|
| **Data collection and processing** | | | |
| Magnification | 29000× | 29000× | 29000× |
| Voltage (kV) | 300 | 300 | 300 |
| Electron exposure ($e^{-}/Å^2$) | 50 | 50 | 50 |
| Defocus range (μm) | -1.2 to -1.5 | -1.2 to -1.5 | -1.2 to -1.5 |
| Pixel size (Å) | 0.97 | 0.97 | 0.97 |
| Symmetry imposed | C3 | C3 | C1 |
| Initial particle (no.) | 76,948 | 68,930 | 52,700 |
| Final particle (no.) | 110,546 | 183,114 | 44,293 |
| Map resolution (Å) | 3.55 | 3.81 | 3.92 |
| Local resolution range (Å) | 2.0-4.8 | 3.2-5.5 | 3.2-5.7 |
| Map sharpening $B$ factor ($Å^2$) | -150 | -150 | -150 |
| **Refinement** | | | |
| R.m.s. deviations | | | |
| Bond lengths (Å) | 0.019 | 0.022 | 0.020 |
| Bond angles (°) | 1.742 | 1.953 | 1.927 |
| Validation | | | |
| MolProbity score | 1.18 | 1.61 | 1.34 |

| | | | |
|---|---|---|---|
| Clashscore | 1.97 | 3.12 | 1.85 |
| EMRinger score | 2.75 | 0.65 | 2.36 |
| Map correlation coefficient | 0.74 | 0.63 | 0.71 |
| Ramachandran plot | | | |
| Favored (%) | 96.63 | 91.39 | 94.07 |
| Allowed (%) | 3.29 | 8.37 | 5.68 |
| Side chain rotamer outlier (%) | 0.08 | 0.24 | 0.25 |
